# Widespread synchronization of codon usage in functionally related genes

**DOI:** 10.64898/2025.12.10.693408

**Authors:** José L. López, Aristeidis Litos, Malte Siemers, Meine D. Boer, Erica Padial, Swapnil P. Doijad, Bas E. Dutilh

## Abstract

The usage of synonymous codons varies along the genome, with strong biases in conserved and highly expressed genes that are optimized for efficient translation. The extent to which selection shapes codon usage in other genes, as well as the associations between gene function, gene expression, and codon usage, remains an important open question. We developed and optimized a novel approach to detect synchronized changes in codon usage patterns and applied it to 15,005 species-representative bacterial genomes spanning the 15 most represented phyla. We show that codon usage is extensively shaped by selection in both highly and lowly expressed genes, with at least ∼20-46% of gene families showing synchronized codon usage evolution across genomes. We reveal that gene pairs with parallel codon usage adaptation are co-expressed, co-regulated, metabolically connected, and functionally associated. By identifying synchronized codon usage evolution between gene pairs, we have generated a genome-wide set of functional associations reflecting correlated expression across species. This underappreciated layer of coordinated codon usage adaptation has important implications for function discovery and engineering.

## Introduction

Codons that translate to the same amino acid (synonymous codons) are used at different frequencies between genes depending on mutational and selective forces (Sharp et al. 1993; Plotkin and Kudla 2010). Current models of codon usage (CU) evolution propose that genomes reach a steady-state in which their gene codon usage is at a selection-mutation-drift equilibrium, where highly expressed genes are predominantly impacted by selection, whereas codon usage in lowly expressed genes is shaped mostly by mutational biases and drift (Bulmer 1991; Bénitière et al. 2025; Cope and Shah 2025). Since highly expressed genes represent only a small portion of protein-coding sequences in a genome, mutation bias and drift are therefore considered to shape codon usage in most genes, and in the genome overall (Cope and Shah 2025).

However, the relative importance of neutral and selective factors in shaping CU is debated (Kudla et al. 2009, Shah and Gilchrist 2010, Hershberg and Petrov 2010, Plotkin and Kudla 2010), and many selective factors have been identified (Hanson and Coller 2018). Selection for efficient translation has led to a strong codon usage bias (CUB) in ribosomal proteins of fast-growing bacteria (Ikemura 1985). Although initiation, dictated by 5’ RNA stability and the Shine Dalgarno accessibility, is the rate-limiting step of translation (Kudla et al. 2009; Allert et al. 2010; Bentele et al. 2013; Goodman et al. 2013; Cambray et al. 2018; Osterman et al. 2020; Nieuwkoop et al. 2023), codon usage contributes to overall translation efficiency by affecting the rate and accuracy of protein elongation, the availability of ribosomes in the host cell, and the mRNA decay, among other processes (Plotkin and Kudla 2010; Quax et al. 2015; Hanson and Coller 2018). The elongation rate is affected because optimal codons match the more abundant tRNA species in the cell (Ikemura 1985; Qian et al. 2012), meaning that the mRNA may be processed faster, allowing the ribosome to be freed to process other transcripts. Thus, an adequate pool of free ribosomes can be maintained, improving overall translation (Shah et al. 2013; Frumkin et al. 2018). Codon usage may also affect translation accuracy, reducing the costs of both missense and nonsense errors (Akashi 1994). In line with this, codon usage bias is higher in longer and more conserved genes (Stoletzki and Eyre-Walker 2007; López et. al 2020). Additionally, codon usage affects mRNA stability and decay (Presnyak et al., 2015, Wu et al. 2019). For example, in yeast and humans, this process involves the DEAD-box helicase Dhh1 (Radhakrishnan et al., 2016) and the protein DHX29 (Hia et al., 2026), both of which sense suboptimal codons and promote initiation of mRNA decay.

Codon usage also reflects fine-tuning of expression across different physiological conditions (Elf et al. 2003; Najafabadi et al 2009; Xu et al. 2013; Zhou et al. 2013), and CUB is correlated with subunit stoichiometry in polycistronic genes encoding protein complexes, showing protein demands of genes tightly condition their codon usage (Quax et al. 2013). While CU has been shown to correlate in functionally or metabolically related genes (Fraser et al. 2004; Lithwick 2005), these studies were limited in scope, focusing on relatively few functions and organisms, with their implication for functional discovery being largely overlooked. Thus, the breadth of these correlated codon usage patterns across functional categories and throughout the bacterial domain remains largely unexplored. Here, we found that codon usage fine-tuning reflects expression requirements and that this signature is extensively present across pairs of co-expressed gene families as they evolve in different bacterial lineages. To test this, we used comparative genomics of thousands of bacterial genomes to investigate the link between protein function and codon usage adaptation. By analysing CUB patterns between pairs of genes, we show that parallel CU adaptation across genomes is strongly shaped by selection in functionally related genes with different expression levels, and that its codon usage synchronization extends across gene families and phyla, revealing a finely tuned code that reflects gene expression patterns and can be leveraged for protein function discovery and a better understanding of bacterial physiology.

## Results

### Codon usage bias is broadly conserved within gene families

Previous research revealed a spectrum of CUB values ranging from highly expressed, conserved genes to lowly expressed, accessory ones (Davis and Olsen 2010; López et al. 2019; López et al. 2020). As CUB can be calculated in different ways, we first benchmarked seven commonly used CUB indices and found that gene codon bias (GCB) best correlated with expression data under a range of conditions, where highly expressed genes with optimized codon usage have high GCB values (Merkl 2003; Sastry et al. 2019; Sastry et al. 2024) (Supplementary Fig. 1 and Supplementary Fig. 2, Supplementary Note 1). Next, we analyzed genome sequences from 2,978 representative species in the five most abundant bacterial phyla, including *Actinomycetota*, *Bacillota*, *Bacillota A*, *Bacteroidota*, and *Pseudomonadota* (Supplementary Table 1). We used GCB to calculate CUB in all encoded gene families and assessed the CUB per functional category (Fig. 1a, b). As expected, genes involved in protein processing and energy metabolism are generally highly optimized, categories that include critical functions for fast-growing bacteria (Karlin et al. 2001, Vieira-Silva et al. 2011). In contrast, unannotated genes were the least optimized (Fig. 1a), illustrating annotation bias: widespread, highly expressed genes are more likely to be studied and thus functionally annotated (Grigson et al. 2025). However, most broad functional categories do not show a clear pattern, likely reflecting that individual functions within each category require different expression levels. To test if orthologs have similar CUB values in different bacteria, we compared single-copy gene families (PGFams) shared between genomes and observed consistent overall GCB patterns (Fig. 1c). Correlation values were lower for the subset of unannotated proteins than for annotated ones (Fig. 1d), in line with the higher expression, conservation, and annotation rates of core genes, and the evolutionary variability of accessory, often hypothetical, genes (Davis and Olsen 2011; López et al. 2020).

**Fig. 1.**
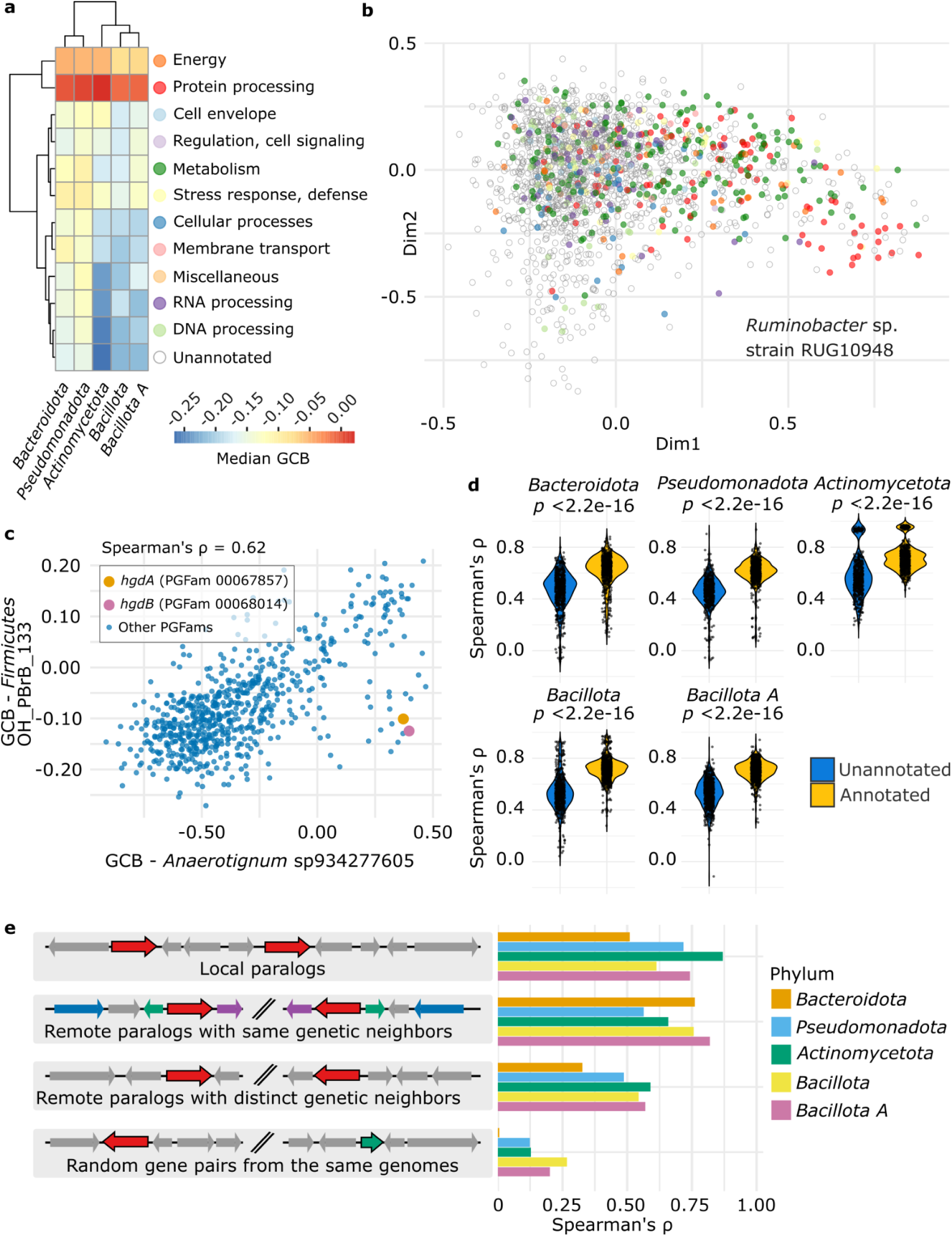
Overall codon usage optimization across bacterial functions. **a** Median GCB values of genes by functional category (Overbeek et al. 2013). The heatmap represents the median of the per-genome median GCB values for each phylum. **b** Example case showing the first two dimensions of a factorial correspondence analysis of per-gene codon usage vectors in *Ruminobacter* strain RUG10948 (*Pseudomonadota*). Genes are colored by functional category as in panel A. **c** CUB values of shared orthologous genes between *Anaerotignum sp934277605* and *Firmicutes* strain OH_PBrB_133 are positively correlated (Spearman’s ⍴ = 0.62). The *hdgA* and *hdgB* genes are highlighted, these are further analyzed in Fig. 2. **d** Spearman correlations between GCB values of orthologous genes (single-copy PGFams) between genome pairs. Violin plots are split by annotation status of the genes. Statistics were performed with Wilcoxon rank-sum tests. **e** Spearman correlations between the CUB values of paralogs (PGFams with two copies per genome, highlighted red) across phyla (Supplementary Table 2 and Supplementary Note 1). Genetic neighbors are defined as PGFams within ±20 genes up- or downstream of a target gene. Local paralogs have <40 genes between them, others are remote. Random gene pairs show weak genome-wide CUB correlations.

We then explored CUB patterns in paralogous genes within the same genome (Supplementary Note 2) and found that correlation was stronger in colocalized paralogs within the same genomic region, as well as in remote paralogs with similar neighboring genes, compared to remote paralogs without shared neighbors (Fig. 1e, Supplementary Table 2, and Supplementary Fig. 3). Together, these results show that CUB is conserved between orthologs, likely reflecting similar expression levels within and across genomes.

### Enzymatic subunits show synchronized codon usage adaptation

Next, we used our genome collection to investigate potential codon usage (CU) correlations between functionally related genes. As an example, we focus on *hgdA* and *hgdB*, which encode two subunits of 4-hydroxybenzoyl-CoA reductase, an oxygen-sensitive enzyme involved in the anaerobic degradation of aromatic compounds (Glöckler et al. 1989). As shown above (Fig. 1c), PGFams generally had similar CUB values in different genomes, e.g. *Firmicutes* strain OH_PBrB_133 and *Anaerotignum* sp934277605, but *hdgA* and *hdgB* formed an exception, showing increased CUB in *Anaerotignum* sp934277605 (Fig. 1c, and Fig. 2a, b).

**Fig. 2.**
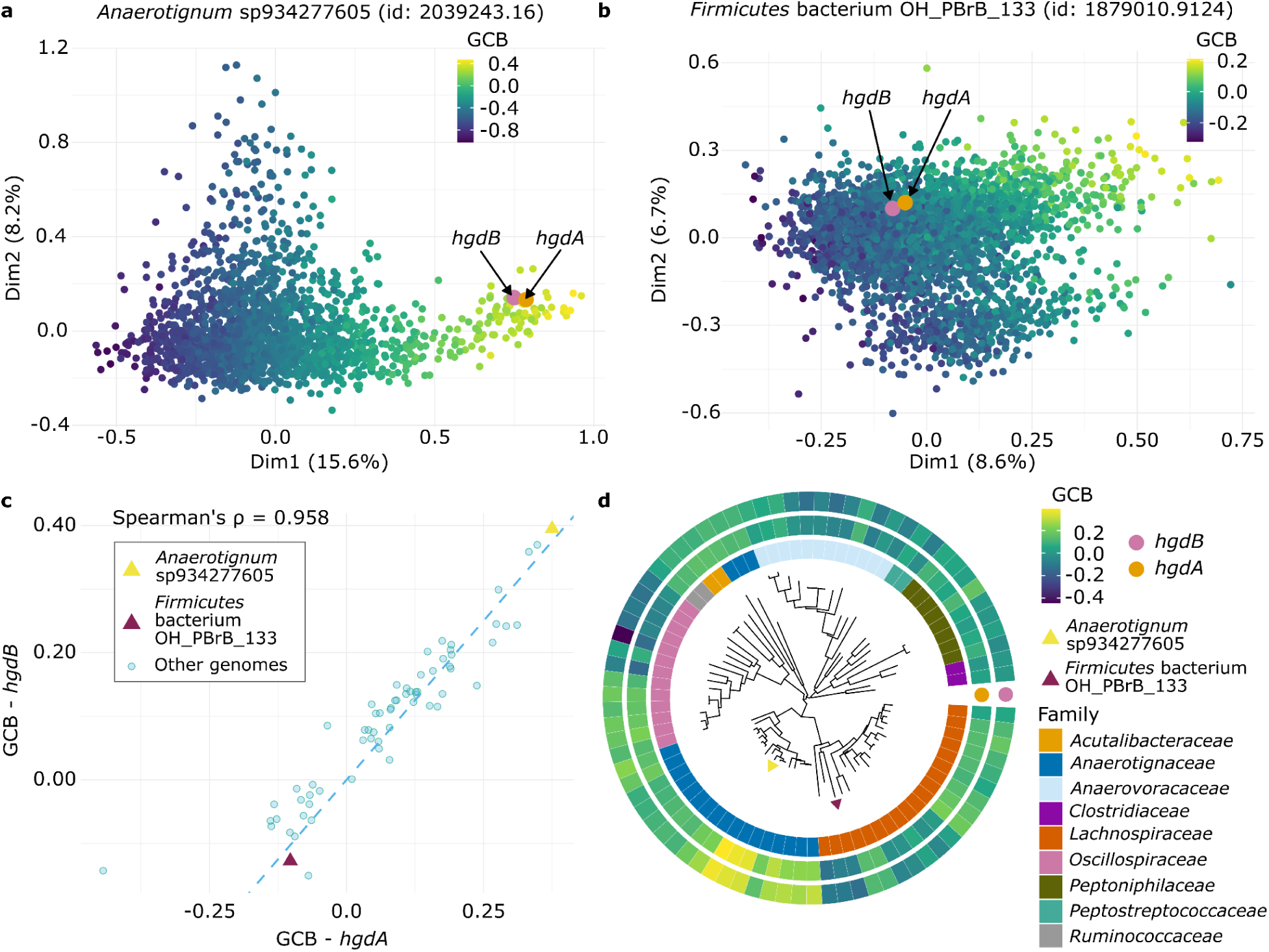
CUB patterns in subunits of 4-hydroxybenzoyl-CoA reductase. **a, b** First two dimensions of a factorial correspondence analysis of per-gene codon usage vectors for two *Bacillota A* genomes: **a** *Anaerotignum* sp934277605 and **b** *Firmicutes* strain OH_PBrB_133. Genes are colored according to their GCB values, with *hdgA* and *hdgB* genes being highly optimized in *Anaerotignum* sp934277605. **c** GCB values of *hdgA* and *hdgB* genes across reference species in *Bacillota A* are strongly correlated (Spearman’s ⍴ = 0.958), indicating co-optimization of the enzyme subunits across genomes. The genomes shown in panels **a** and **b** are indicated. **d** The phylogenetic tree of genomes from *Bacillota A* coding for *hdgA* and *hdgB* was subsetted from the bac120_r220.tree, obtained from GTDB (Parks et al. 2021). Color strips indicate taxonomic families and GCB values, example genomes are highlighted.

Across species, we observed strong CU synchronization between these genes, with *Firmicutes* OH_PBrB_133 and *Anaerotignum* sp934277605 at opposite ends of the optimization spectrum (Spearman’s ⍴ = 0.958, *p* < 2.2e-16, Fig. 2c), indicative of divergent selection pressures. In *Anaerotignaceae*, which inhabit strictly anaerobic environments, aromatic compound degradation is likely essential for energy metabolism (Brackmann et al. 1993; Gibson et al. 1997; Ueki et al. 2017), explaining the higher CUB in *hgdA*/*hgdB*. In contrast, *Lachnospiraceae*, common gut bacteria that rely primarily on polysaccharide fermentation (Meehan and Beiko 2014; Sorbara et al. 2020), may have reduced expression, reflected in a lower CUB of these genes. In this view, the codon usage of both genes is adapted to their required expression levels, and because they are part of the same enzyme, their similar codon usage reflects their shared co-expression. We therefore refer to this pattern as synchronized codon usage evolution (SCUE).

### Synchronized codon usage evolution (SCUE) across functionally related proteins

To quantify the SCUE between protein-coding genes on a large scale, we correlated GCB and GCB z-scores between all PGFam pairs across genomes (Supplementary Note 1). We hypothesized that additional factors beyond PGFam identity, such as phylogenetic signal (Pagel M 1997), genomic mean GCB, and genomic GC content, could influence our correlation values, given the strong associations observed between individual PGFam CUB and these factors (Supplementary Fig. 4). To assess these potential confounders, we took 15,003 high-quality genomes from the five major phyla (Supplementary Table 1) and constructed phylogenetically informed regression models for each PGFam pair, incorporating the confounders as covariates and considering potential interactions. Based on these SCUE models (Supplementary Table 3), we generated graphs where edges represent pairwise relationships with *t* values between PGFams. We next analysed the functional connections by defining SCUE clusters. First, we applied iterative Louvain clustering (Blondel et al. 2008), and then assessed the STRING association score (Szklarczyk et al. 2022) of PGFam pairs within these clusters as a proxy for known functional interactions (Supplementary Table 3).

We benchmarked the different approaches used here to optimize the mean STRING combined interaction score between gene pairs within the same SCUE cluster. First, the STRING score increased significantly when phylogenetic signal, genomic mean GCB, and GC content were included as covariates (Supplementary Fig. 5, left panels). Furthermore, using normalized GCB z-scores instead of raw GCB values further improved functional linkages (higher mean STRING scores, Supplementary Fig. 5, right panels). Finally, using the *t* values (Phylolm regression estimates normalized by their standard errors) increased the number of detected clusters while maintaining comparable STRING scores (Fig. 3a). Accordingly, we selected the phylogenetic linear models that used GCB z-score as input and incorporated the covariates and interactions; the resulting *t* values were used for all subsequent analyses.

**Fig. 3.**
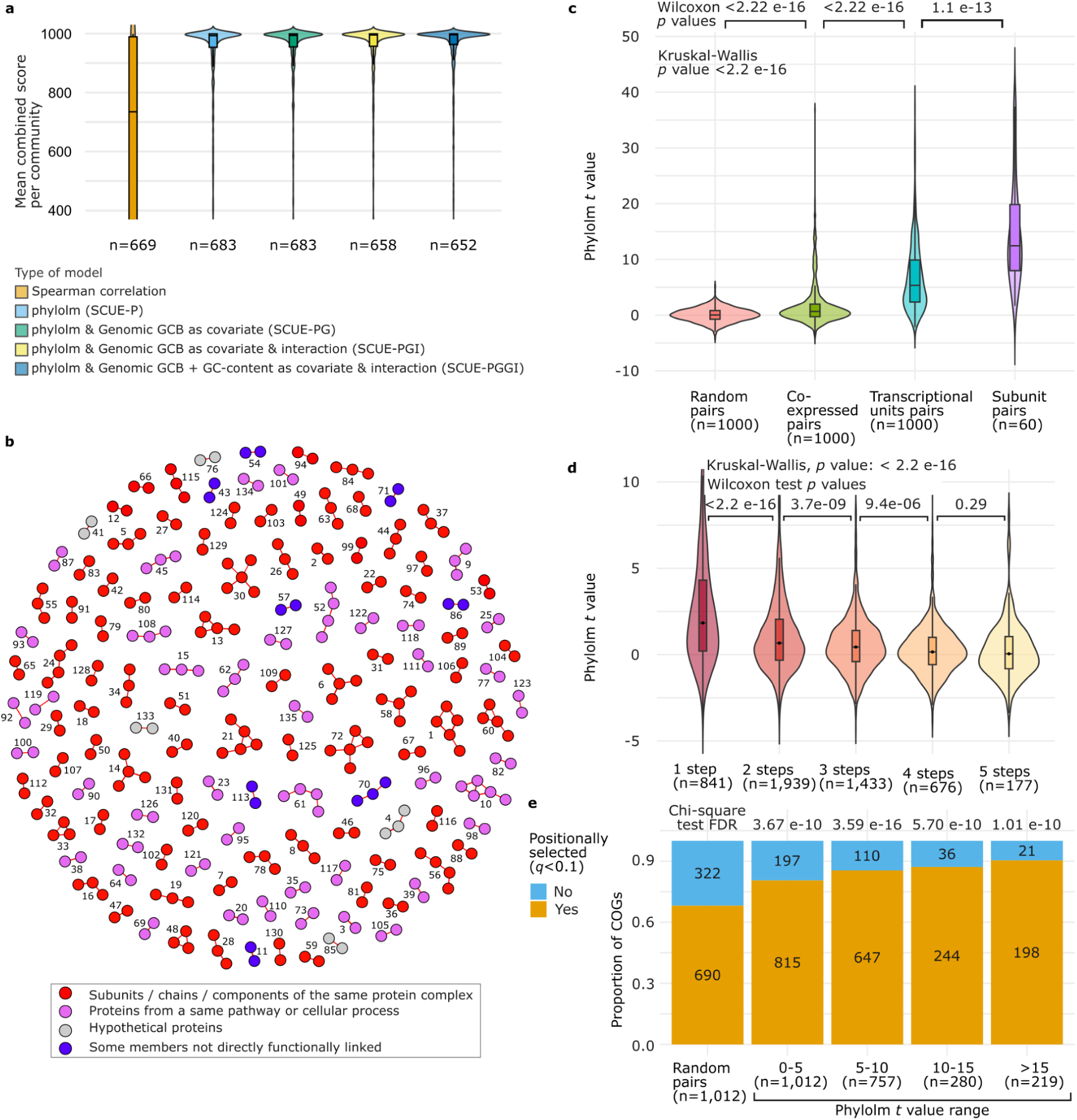
Synchronized CU evolution between PGFam pairs. **a** STRING combined scores for clusters obtained using Spearman’s ⍴ values and *t* values of different SCUE models that account for phylogenetic signal, genomic GCB, and GC content as covariates with interaction terms; n represents the number of clusters with STRING interactions (>2 members). **b** The top 200 positively correlating genes from the optimal *Pseudomonadota* model are visualized as a network. Colors represent different types of functional relatedness, and numbers denote the cluster identifier (Supplementary Table 2). **c** Phylolm *t* value distributions increased from random gene pairs, to co-expressed pairs, to co-transcribed genes, to genes encoding subunits of the same protein complex. iModulons and transcriptional units from *E. coli* K-12 MG1655 were used. **d** Phylolm *t* values from SCUE-PGGI models of enzymes that are closer in metabolic pathways are higher than more distant ones (1-5 steps apart in the metabolic network). **e** Positionally selected functions are more prevalent among pairs with high SCUE-PGGI *t* values. The proportion of positionally selected PGFams in pairs with Phylolm *t* values within ranges: 0-5, 5-10, 10-15, and >15, are compared with random pairs, using chi-squared tests.

These results indicate that genomic factors, major confounders in assessing CU synchronization among genes, can be effectively accounted for by using phylogenetic multiple regression with interaction effects. Using GCB z-scores as input already corrected for biases including genomic GCB, GC content, and potential other unknown confounders. Subsequent correction for the strong phylogenetic signal (Fig. 2d, and Supplementary Fig. 6) further improved model performance (Supplementary Fig. 5). Finally, we compared the GCB index with the more commonly used indices CAI and E (Supplementary Fig. 7). GCB resulted in slightly higher STRING scores, consistent with our initial selection criteria (Fig. S1), while CAI and E yielded more clusters. Overall, these methodological advances allowed us to reveal that codon usage is under selection in a broad array of gene functions.

Subgraphs consisting of the top 200 strongest synchronized PGFam pairs (Bonferroni-adjusted *p* < 0.01, Supplementary Table 2) formed small clusters of 2-5 PGFams, most of which included protein families with strong functional links based on SEED annotations (Overbeek et al. 2013) (Fig. 3b, Supplementary Table 2). These links included subunits of the same enzyme, components of enzyme complexes or cellular machinery (red clusters in Fig. 3b), or enzymes from the same pathway or metabolic process (pink clusters). We also identified several novel SCUE links (blue clusters), for which low or no STRING scores were observed (Supplementary Table 3).

Many clusters contained genes with closely related functions, suggesting that SCUE reflects co-expression of functionally linked genes. To test this hypothesis, we compared SCUE *t* values between random gene pairs with those between enzyme subunits, genes within the same transcriptional unit, and co-expressed genes (Fig. 3c). Subunits of enzymes or protein complexes showed significantly higher SCUE *t* values than genes within the same transcriptional unit (Wilcoxon test *p* value = 1.1e-13, Fig. 3c). Genes from the same transcriptional unit, in turn, showed significantly higher SCUE *t* values than co-expressed pairs, while co-expressed pairs had significantly higher values than random pairs (Wilcoxon test, *p* value < 2.22e-16, Fig. 3c). These results support a model where shared patterns in CU evolution across genomes reflect the co-expression of functionally associated genes. Gene pairs that are co-expressed in only a few genomes will not result in significant SCUE values in our models, which are based on correlations across many genomes. SCUE pairs are also most likely to have correlated expression levels to achieve a strict stoichiometric demand. The weakest association is between PGFams that are co-regulated only in one genome (e.g., *E. coli* in Fig. 3c), which may span multiple transcriptional units (Catoiu et al. 2024). Next, genes within a single transcriptional unit are more tightly co-regulated, co-expressed, and functionally associated across many other genomes, if operon structure is conserved. Finally, this effect is strongest for subunits or components of protein complexes, whose genes require highly coordinated expression to maintain balanced cellular dosage (Papp et al. 2003; Fraser et al. 2004; Lithwick 2005; Quax et al. 2013), and are most likely co-expressed in many species.

To investigate the SCUE between metabolically associated genes, we quantified the distance between enzymes in metabolic networks. Enzymes separated by a single metabolic step showed significantly higher SCUE values (Wilcoxon test, *p* value < 2.22e-16), followed closely by those separated by two and three steps (3.7e-09 and 9.4e-06, respectively, Fig. 3d), although SCUE rapidly declined with increasing metabolic distance. The effect was stronger for enzymes within the same subsystem subclass and was consistently observed across all five phyla (Supplementary Fig. 8). These results suggest that the closest metabolically connected enzymes correlate in their expression across genomes, likely reflecting more dependent metabolic fluxes, whereas more distant enzymes are less constrained.

A recent study reported that two-thirds of functions are positionally selected on the bacterial genome, revealing an evolutionary mechanism that links gene expression to cellular growth rate (Hu 2025). We hypothesized that SCUE pairs may also be enriched in the positionally selected genes, reflecting a stronger selection of their co-expression. To test this, we compared the proportions of positionally selected genes among SCUE pairs across different ranges of *t* values with those of random gene pairs. Indeed, SCUE pairs showed a significant signature of positional selection (Chi-square test, *p* value < 1.01e-10, Fig. 3e), supporting CU adaptation and positional selection as complementary evolutionary layers that reflect fine-tuned gene expression.

Additionally, significant gene pairs spanned the genome, with 47% of pairs being close by (i.e., separated by 5 genes or fewer) and 53% consisting of functionally related genes located far apart in the genome (>5 genes apart, Supplementary Figure 9), which shows that SCUE reflects correlated expression also occurring between functionally related genes located far apart in the genome.

### SCUE in half of the protein families across bacterial phyla

Above, we focused on the five most species-rich phyla because they contain the most genomes in the database, and thus likely contain the strongest SCUE signal. To generalize these observations, we extended our models to include ten phyla with >100 available representative species genome sequences, including *Acidobacteriota*, *Bacillota C*, *Bdellovibrionota*, *Campylobacterota*, *Chloroflexota*, *Cyanobacteriota*, *Desulfobacterota*, *Planctomycetota*, *Spirochaetota*, and *Verrucomicrobiota*. The SCUE pairs identified within these additional phyla had clear functional connections, although fewer were detected within these additional phyla (Supplementary Fig. 10, and Supplementary Table 4). Thus, SCUE is widespread in the bacterial domain. Using stringent multiple-testing correction (Bonferroni *p* < 0.01), approximately 77% of gene pairs with significant SCUE also had high STRING combined scores (>700, Supplementary Table 4). For the remainder of cases, i.e. 23% of the gene families (n=1,051), this suggests that SCUE provides new evidence about their function.

With the stringent Bonferroni criterion used above, only 20% of all PGFams had at least one significant SCUE partner (Supplementary Table 4). To reduce the stringency of multiple-testing correction and reveal more functional associations, we applied a Benjamini-Hochberg (BH) procedure. To identify a good cutoff, we selected SCUE pairs at a range of different BH cutoff values, and determined when the median STRING combined score of the significant pairs exceeded 700 (Supplementary Fig. 11, and Supplementary Table 4). Using this approach, we found that approximately 46% of all PGFams (ranging from 14% to 60% for individual phyla) had at least one significant SCUE partner, and about 30% of all PGFams (ranging from 8% to 43% across phyla) had a significantly SCUE partner with a high STRING interaction score (combined score >700, Supplementary Table 4). These results indicate that CU is under significant adaptive selection in a substantial fraction of bacterial functions. The remaining PGFams without a detected SCUE partner may correspond to genes with more variation in CUB values across genomes, suggesting that alternative modelling strategies may be necessary to capture their SCUE signal. Alternatively, these genes may not co-express with others, e.g. because they represent independently functioning enzymes.

### Synchronized gene pairs have similar CUB values, codons, and amino acids

Above, we tested whether gene pairs had correlating CUB patterns across genomes. To test whether they also have similar CUB values, we calculated the median absolute difference in GCB z-scores (|ΔGCBz|) within their genomes, and compared these values between non-significantly and significantly SCUE pairs (Supplementary Fig. 12a). The latter gene pairs typically fell within the lowest 5-10% of median |ΔGCBz| values for pairs of a given PGFam and had significantly lower median |ΔGCBz| values, indicating that synchronized genes tend to have similar codon usage bias (Supplementary Fig. 12b, c, d). These results reveal that SCUE gene pairs closely track one another as their CU evolves across genomes, and suggest that median |ΔGCBz| could serve as an additional criterion to enhance detection of functionally linked pairs.

To determine whether similar CUB values reflect similar codon composition, we calculated codon composition distances between gene pairs corresponding to all PGFams included in our models. We found that SCUE pairs had significantly smaller codon composition distances than non-significant pairs (Supplementary Fig. 13a), consistent with their more similar CUB values (Supplementary Fig. 12). Interestingly, SCUE pairs also had significantly similar amino acid composition (Supplementary Fig. 13b). This not only contributes to their shared codon composition, but might also reflect an additional layer of selection. Indeed, the positive correlation between codon and amino acid distances was stronger in significant than in non-significant pairs (Spearman’s ⍴ values of 0.78 and 0.62, respectively, Supplementary Fig. 13c). Mirroring the patterns observed for median |ΔGCBz| values, the most significant SCUE pairs typically fell within the lowest 10–15% of codon composition distances and the lowest 15–20% of amino acid composition distances (Supplementary Fig. 13d). This suggests an evolutionary coupling between codon and amino acid compositions in SCUE pairs where both might reflect protein expression optimization.

### Synchronized codon usage evolution is ubiquitous across expression levels and bacteria

To explore how SCUE varies with different gene expression levels, we used |ΔGCBz| values and codon usage distances in significant pairs across expression levels, using either experimental data, or GCB as a proxy for expression. CU models of selection-mutation-drift posit that highly expressed genes present a higher impact of selection over mutational bias compared to lowly expressed genes (Shah and Gilchrist 2011). If selection on codon usage is stronger in highly than in lowly expressed genes, we would expect a negative correlation between |ΔGCBz| and expression values, as mutation and drift would mask weak selection for codon usage bias in lowly expressed pairs. Overall, neither |ΔGCBz| nor codon distance values between gene pairs correlate with expression (Supplementary Fig. 14a-g), although we do observe a majority of weak negative correlations when analyzing each gene pair individually across genomes (Supplementary Fig. 14h). Thus, these results align with currently accepted models but also show that selection strongly shapes gene codon usages in the lowly expressed genes. These results are consistent with and add to the evidence that previously showed selection impacts both highly and lowly expressed genes (Yannai et. al 2018), and also explain why our models are able to detect SCUE links across all kinds of gene functions, where both highly and lowly expressed gene pairs would contribute to a consistent signal across genomes that is effectively captured by our approach.

Moreover, SCUE models were obtained using representative species that spanned a broad range of lifestyles and growth rates. We estimated predicted minimal doubling times (Weissman et al., 2021) on genomes and observed that |ΔGCBz| values in significant pairs were slightly positively correlated with predicted minimal doubling times (Supplementary Fig. 15), also consistent with mutation-selection-drift models which predict slow growing bacteria have a lower impact of selection shaping codon usage. Altogether, SCUE links provide novel evidence with implications for codon usage evolutionary models, highlighting the need to consider a broader impact of selection on codon usage.

### SCUE identifies candidate novel functional interactions

SCUE complements existing evidence of functional interactions between genes and can be leveraged to identify novel functional partners, prioritising targets for experimental validation (Supplementary Tables 3 and 4). We hypothesize that these novel candidates may involve yet uncharacterized functional links that exhibit physical interactions in the cell, particularly in less-studied bacterial phyla, or functions that require a tight correlated expression across a substantial fraction of the genomes analyzed. Although not physically interacting, the latter cases might represent either indirect functional links between related processes (i.e., metabolically connected biological processes) or unrelated processes that share cofactors, energy dependencies, or unknown metabolites. The high proportion of high-confidence STRING links found among significant SCUE pairs suggests that novel SCUE links represent new functionally linked candidates that will require further experimental validation.

We will present four cases where proteins were not previously recognized to interact. First, we will present the computational evidence, and then summarize experimental results that provide additional support for their functional association.

First, the histone-like *Bacillus subtilis* protein HBsu, an essential DNA-binding protein linked to replication initiation in *B. subtilis* (Micka et al. 1991; Karaboja and Wang 2022), strongly clusters with ribosomal proteins based on CU synchronization (Supplementary Table 3), with high SCUE *t* values and low median |ΔGCBz| values (Fig. 4a, b, and Supplementary Fig. 16a, b) in *Bacillota* (cluster 14, Supplementary Table 3) and *Bacillota A* (cluster 137, Supplementary Table 3) phyla. As STRING currently provides limited evidence for interactions between HBsu and ribosomal proteins, mainly through co-expression and genomic neighbourhood (only two ribosomal proteins have a combined score >700, Supplementary Table 3, and Fig. 4b), our findings point to a complementary role of HBsu and ribosomes in optimizing cell division. This is further supported by regulatory links between HBsu mRNA stability and ribosomal gene expression (Braun et al. 2017), and evidence of a role in chromosome organization (Dame 2005). A similar SCUE with ribosomal proteins (cluster 550, Supplementary Table 3) could be detected for DNA-binding protein HUβ (i.e., Heat-unstable beta subunit) in *Pseudomonadota*, which belongs to the same class of histone-like protein families with similar roles binding DNA (Micka et al. 1991; Hammel et al. 2016). We hypothesize that these proteins could have a role in coupling DNA replication and translation, where HBsu and HUβ optimization may be required to sustain fast growth.

**Fig. 4.**
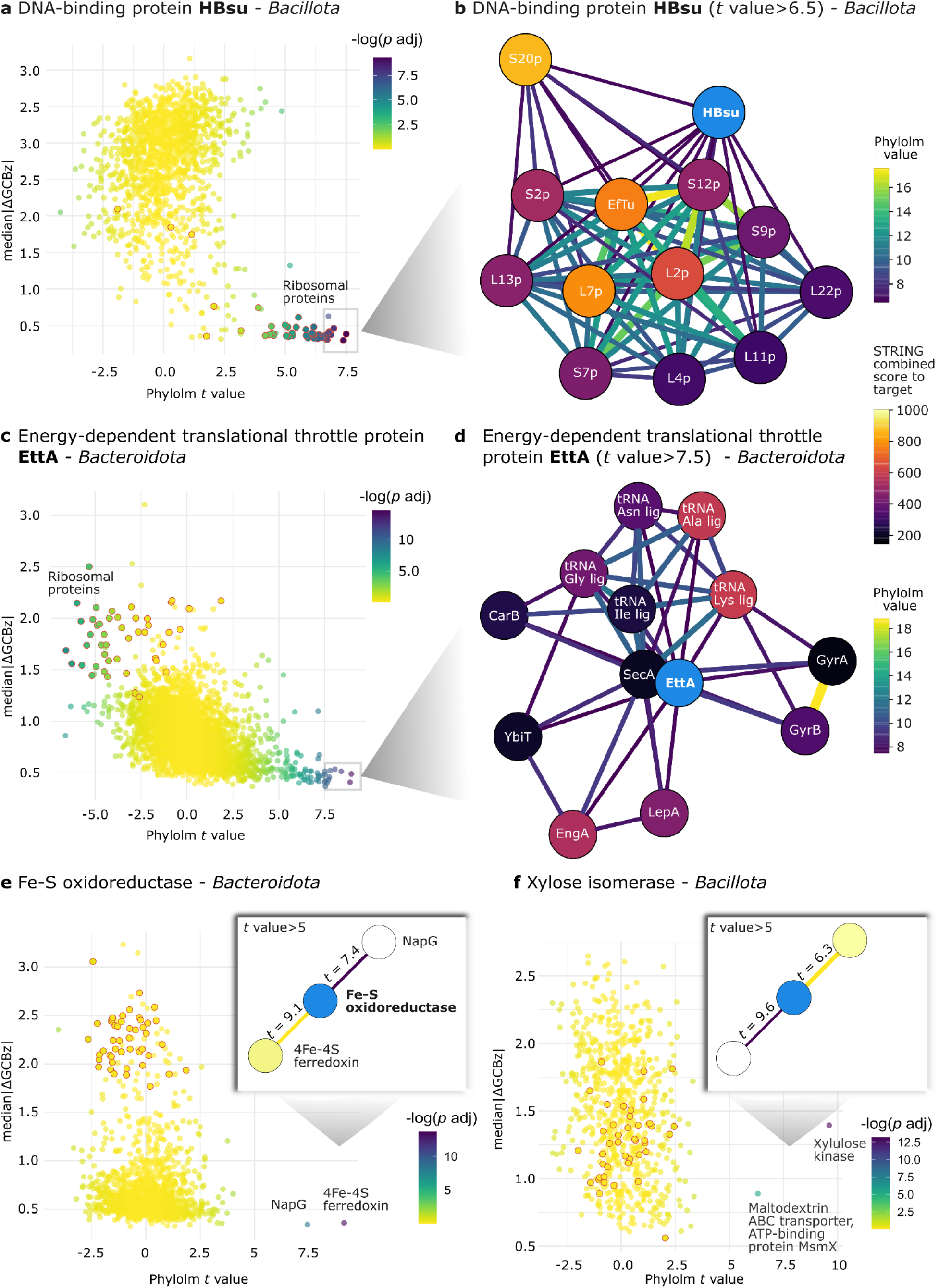
SCUE partners represent known and putative functional interactions. **a, c, e, f** Phylolm *t* values and median |ΔGCBz| values to the target PGFam are shown to represent the space of interactions with all other PGFam pairs. Color indicates the significance of SCUE according to the SCUE-PGGI model. Ribosomal proteins are outlined in red. **b, d** Subsets of significantly SCUE PGFams pairs are represented by nodes with their color representing the highest known STRING combined score in interactions with the target PGFam (in light blue). Edge thickness is proportional to the Phylolm *t* values. Target PGFam and their SCUE partners include (**a** and **b**) DNA-binding protein HBsu, **c, d** energy-dependent translational throttle protein EttA, **e** Fe-S oxidoreductase, and **f** xylose isomerase.

Whereas the example above is closely associated with ribosomal proteins and has high CUB and expression in fast-growing bacteria, there are also interesting examples of the converse. We found that in *Bacteroidota*, CU of the energy-dependent translational throttle protein EttA, an ATPase from the ATP-binding cassette subfamily F (ABC-F) that regulates the translation elongation cycle (Boël et al. 2014), shows significant SCUE with a series of proteins involved in different translation-related functions (all *t* values > 4.5, FDR < 8 e-04, Fig. 4c, d, Supplementary Fig. 16c, d, and Supplementary Table 5). Interestingly, most of these proteins consume either ATP or GTP and are involved in ribosome functioning, yet have no strong known interactions with EttA (low STRING combined scores with weak evidence of co-expression in *Bacteroidota*, Supplementary Table 5). By binding to the ribosome, EttA restricts ribosome entry into the translation elongation cycle at a high ADP:ATP ratio (Boël et al. 2014, Chen et al. 2014). EttA restriction on translation is reflected by its negative *t* values with ribosomal proteins (Fig. 4c, and Supplementary Table 5), suggesting that slow-growing bacteria (i.e., those with lower CUB in their ribosomal proteins) evolved a higher CUB for this protein. Although the physiological function of EttA has not been fully uncovered, we hypothesize that it has a critical role in modulating ribosomal function, especially in slow-growing bacteria that thrive in low nutrient conditions and must tightly couple translation to energy availability. This hypothesis is consistent with the fact that co-expression evidence of EttA with ribosomal proteins is only found in slow-growing bacteria (Supplementary Table 5), and with the loss of EttA ortholog in a beetle symbiont, *Burkholderia* sp. Lv-StB, which may reflect an adaptation to a more steady nutritional niche (Waterworth et al. 2020). Moreover, EttA protects against antibiotics in the slow-growing *Mycobacterium tuberculosis* (Gupta et al. 2010; Cui et al. 2022; Li et al. 2022) but not in fast-growing bacteria such as *Staphylococcus aureus* (Meir et al. 2020) or *Escherichia coli* (Boël et al. 2014), where it has a role in entering the stationary phase (Boël et al. 2014; Chen et al. 2014). Thus, this suggests EttA may be more essential in slow than in fast growers and could hold potential for targeting slow-growing pathogens.

Other SCUE pairs showed no prior evidence of known functional interaction (STRING interactions are absent across genomes). In these cases, SCUE suggests previously unrecognized interactions based on shared components or processes. For instance, cluster 242 in *Bacteroidota* (Fig. 4e, and Supplementary Table 5) includes Fe-S oxidoreductase and 4Fe-4S ferredoxin iron-sulfur binding domain proteins, which are known to interact (STRING combined score = 972) but synchronize its CU with a ferredoxin-type protein NapG (periplasmic nitrate reductase, Fig. 4e), with absent STRING interactions. In *Pseudomonadota*, NapG is involved in transferring electrons from quinol molecules to the nitrate reductase during anaerobiosis, together with other proteins encoded in the *nap* operon (Brondijk et al. 2004). In *Bacteroidota*, only NapG ortholog is present and, although this electron shuttling function is known to interact with other oxidoreductases such as NADH-ubiquinone oxidoreductase (Supplementary Table 5), the Fe-S oxidoreductase (PGF_00004164), which is a SCUE pair, might be an unknown electron shuttling partner, or alternatively they synchronize their CU because they both utilize 4Fe-4S clusters, typical of bacterial ferredoxins (Sparacino-Watkins et al. 2014).

Another SCUE pair brings together xylose isomerase (EC 5.3.1.5), xylulose kinase (EC 2.7.1.17), and the maltodextrin ABC transporter ATP-binding protein MsmX (Fig. 4f, cluster 86 in *Bacillota*). Although MsmX is connected with many other sugar transporters (Supplementary Table 5), its specific connection to xylose is not known at STRING so we speculate a role in transporting specifically this sugar.

We provide all the putative novel interactions (Supplementary Table 5) and a database of PGFam pairs displaying SCUE, along with STRING combined scores and median |ΔGCBz| values for the phyla analyzed here, at https://coadaptation.shinyapps.io/CUCA_PGGI/.

## Discussion

In this study, we implemented phylogenetically aware models to investigate synchronized codon usage evolution (SCUE) in gene pairs and showed that the vast majority of significant SCUE pairs are functionally linked, spanning diverse functions across the genome. Significant SCUE pairs comprise both highly and lowly expressed genes, and neighbouring genes within operons as well as genes located far apart in the genome. Our efforts to optimize the synchronization metrics have provided a vast improvement in sensitivity over early studies that used Spearman correlation (Fraser et al. 2004; Lithwick 2005), and provide a comprehensive view of codon usage similarity throughout the bacterial domain.

There are several widely proposed hypotheses to explain selection-driven changes in codon usage and the emergence of SCUE patterns. First, changes in promoter strength or mRNA accessibility can alter mRNA levels or translation initiation rates, respectively, generating selection pressure that favors codons with higher elongation rates to reduce ribosome occupancy (Kudla et al. 2009, Plotkin and Kudla 2010). Second, changes in the tRNA pool due to increased expression or variation in the copy number of a specific tRNA species (Cope and Shah 2025) can impose selection pressure across all genes recognized by that tRNA. However, as changes in tRNA pools affect many unrelated genes in parallel, SCUE may be better explained by changes in expression level.

In line with a previous report in *E. coli* (Yannai et. al 2018), our observation that SCUE pairs ubiquitously include both low- and high-expressed genes has implications for models of codon usage evolution (Bulmer 1991; Bénitière et al. 2025; Cope and Shah 2025). These models postulate that mutational biases and drift have the greatest impact on codon usage in lowly expressed genes, in slow-growing bacteria, and, more broadly, across most genes in a genome, while codon usage in highly expressed genes is mostly shaped by selection. Although our results are consistent with these factors having a somewhat greater impact on lowly than on highly expressed genes (Supplementary Figure 14h), the presence of widespread CU synchronization, even among lowly expressed genes and diverse functions, suggests that selection plays a more substantial role than previously thought. Selective factors that could play an important role include mRNA stability and decay, which have been linked to codon optimization and to functionally related genes in eukaryotes (Presnyak et al. 2015, Wu et al. 2019, Zhang et al., 2025). Indeed, correlated expression is influenced both at the level of protein synthesis and mRNA decay.

Our analysis of gene pairs detected SCUE in at least half of the bacterial functions by their CU patterns across hundreds of species. However, selection likely also shapes codon usage in less ubiquitous genes, or in autonomous genes that do not require partners for proper functioning. We expect that both additional sampling (newly sequenced bacteria) and potential extensions to our current approach will further strengthen the detection of codon usage adaptations, facilitating its association to functions or phenotypes.

As SCUE affects genes with both high and low expression levels, our metric is a valuable addition to the functional association toolbox. Indeed, significant SCUE pairs are functionally related, many already known (high STRING scores), but also promising novel candidates. Thus, SCUE represents a novel approach for identifying functional dependencies, and we provide a comprehensive resource of SCUE links including both known and previously uncharacterized functional connections.

Significant SCUE involves gene pairs that exhibit correlated CU across hundreds of genomes and likely represent gene pairs that are subject to strict stoichiometric dependency (e.g., subunits in protein complexes), flux balance constraints in metabolism (e.g., connected enzymes in pathways), or other linked metabolic processes. Beyond functional discovery, SCUE links may contribute to functionally characterizing which genes within the same functional co-expression module require tight co-expression. Further experimental validation of these correlations could open the way to broadly use CU to describe bacterial lifestyle and physiology, with potential applicability in metagenomics.

Detecting SCUE pairs across the Tree of Life will enhance protein-protein interaction predictions, provide greater insight into known functional partners, reveal novel indirect functional links connecting related processes, and help assign new functional annotations to otherwise uncharacterized proteins. This opens up the possibility of applying our models to the discovery of novel biosynthetic gene clusters, anti-defense systems, and other uncharacterized functional modules. We anticipate that these patterns will be valuable for metabolic modeling and engineering, because codon optimization reflects gene expression and may therefore also capture, to some extent, proteome partitioning and potentially metabolic fluxes. Additionally, CU signatures may aid in predicting biome or environmental preferences and in broadly describing life-history strategies across bacteria, archaea, and eukaryotes. Overall, we reveal at an unprecedented scope and scale that codon usage is under strong selective pressure, likely fine-tuned to match gene expression levels across a broad range of functions throughout the bacterial domain.

## Materials and Methods

### Codon usage bias index selection

We calculated and compared different codon usage bias (CUB) indices to expression data from the Precise 1k dataset from *Escherichia coli* K-12 MG1655 (Sastry et al. 2019) and the MODULOME dataset from *Bacillus subtilis* 168 (Sastry et al. 2024). Expression data consisted of per-sample log(TPM)-normalized values from selected RNA-seq samples used by the authors in the iModulon pipeline (Sastry et al. 2024).

CUB indices included CAI (Sharp and Li 1987), MELP (Supek and Vlahoviček 2005), E (Karlin, S. and Mrázek 2000), GCB (Merkl 2003), MCB (Urrutia and Hurst 2001), SCUO (Wan et al. 2004), ENC (Wright 1990), and Fop (Ikemura 1981), and were computed using the coRdon R package (Elek 2019). As input, we used nucleotide coding sequences and a list of putatively highly expressed genes, either genes encoding ribosomal proteins or a set of manually selected highly expressed genes (PHX) based on proteome data from the PaxDB database (Wang et al. 2015). Proteins shorter than 80 amino acids were excluded, as codon bias measures are generally unreliable for short sequences.

To evaluate each index’s ability to predict gene expression, we calculated Spearman correlations between each index and the expression data. To avoid biases introduced in the original GCB index (see Supplementary Notes), we included a custom modified version of GCB (GCB mod), consisting of the initial GCB calculation but omitting the subsequent optimization loop. While GCB only allowed us to include a subset of genomes, GCB mod enabled the inclusion of all high-quality genomes in our analyses. We used the GCB index on a smaller subset of genomes for the first part of our analyses, including the results shown in Fig. 1, and the GCB mod index on all genomes (Supplementary Table 1) for the remaining analyses in this study, both using ribosomal protein-coding genes as the reference set. For simplicity, we refer to GCB mod as to GCB in our models and analyses from Fig. 2 onwards.

To normalize CUB values within each genome and reduce the impact of genomic confounders, we calculated GCB z-scores by subtracting the genome-wide mean GCB and dividing by its standard deviation.

### Genome selection and annotation

We downloaded genomes from the BV-BRC database (Olson et al. 2023; https://www.bv-brc.org/) as of September 2023 that were also listed as reference species in GTDB release 207 (Parks et al. 2021). These genomes were affiliated with the five main phyla, *Bacillota*, *Bacillota A*, *Actinomycetota*, *Pseudomonadota*, and *Bacteroidota*, as well as an additional 10 less-represented phyla (Supplementary Table 1). We retained genomes meeting the following criteria: more than 30 ribosomal proteins, more than 1,000 CDS, and CheckM (Parks et al. 2015) completeness and contamination scores above 80% and below 10%, respectively. Finally, we only retained genomes for which ribosomal protein genes had significantly lower Nc value (Mann-Whitney test *p* value<0.01) than the rest of the genes in the genome, to avoid the influence of other dominant selective pressures, such as GC content and G+C strand bias, following the rationale of Lithwick *et al*. (2005).

### Functional annotations from the BV-BRC database

We used the “features.tab”, “pathway.tab”, and “subsystem.tab” annotation files from the BV-BRC database, which provide SEED-based annotations of protein functions and their associations with specific metabolic pathways, subsystems, subclasses, and classes (Overbeek et al. 2013). Cross-genus protein families (PGFams) were used to group orthologous protein-coding genes, offering a standardized approach to represent and compare the protein content of bacterial genomes in BV-BRC (Davis et al. 2016).

### Genomic mean GCB calculation

The mean GCB of all genes in a genome, here referred to as the genomic mean GCB, reflects the strength of selection on codon usage. Genomes with lower genomic mean GCB exhibit stronger selection, as ribosomal genes are more distinct from the rest of the genes, resulting in most genes having more negative values (Merkl 2003).

We obtained GC content of genomes from GTDB-associated metadata. To reduce collinearity between covariates and interaction terms, both mean GCB and GC content were centered by subtracting their mean across genomes and used as co-variates in the SCUE models.

### Synchronized codon usage evolution (SCUE) in PGFam pairs across genomes

For genes belonging to single-copy PGFams per genome, we calculated pairwise Spearman correlations, as well as a series of phylogenetically informed regression models (SCUE models, described below), using GCB, GCB z-scores, CAI, and E as CUB input indices for benchmarking. For the five main phyla, only PGFam pairs that co-occurred in at least 100 genomes were included, ensuring sufficient power to detect significant correlations across species. For the additional phyla, this threshold was reduced to 50. PGFams with more than one representative gene per genome were excluded. Since these correlations reflect CUB between pairs of genes, significantly positive values are considered SCUE.

SCUE models were implemented using the phylolm R package and a GTDB-based phylogenomic tree (bac120_r220.tree) subsetted to include the selected genomes for each phylum. Controlling the phylogenetic signal allows us to correct regression models for CU patterns driven by shared ancestry rather than true codon usage synchronization reflecting co-expression between gene pairs (Pagel M 1997).

For each pair, CUB values of one PGFam were used to model the values of the other. Four models were fitted, all incorporating the phylogenomic tree: 1) a model controlling only for phylogenetic signal (SCUE-P); 2) a model including genomic mean GCB as a co-variate (SCUE-PG); 3) a model including genomic mean GCB as a covariate with an interaction term with the independent variable (SCUE-PGI); and 4) a full model including genomic mean GCB and genomic GC content as covariates, each with interaction terms (SCUE-PGGI). Model estimates and *t* values (estimate / SE) were then extracted to quantify codon usage synchronization for each PGFam pair (Supplementary Table 3).

### iModulon, transcriptional units, subunits, and metabolic analyses

We obtained iModulon tables from *Escherichia coli* K-12 MG1655 (Catoiu et al. 2024) and mapped the genes within each iModulon to their corresponding PGFam annotations, thereby grouping PGFams that were part of the same iModulon. Genes belonging to the same transcriptional units in *Escherichia coli* K-12 MG1655 were obtained from EcoCyc, and their corresponding PGFam annotations were mapped to identify PGFams belonging to the same transcriptional unit. Subunits of the same enzyme were identified in the RAST-based functional annotations by searching the keywords “subunit” and “chain”, followed by manual verification. PGFams with Enzyme Commission (EC) annotations were selected within each phylum to assess metabolic connections. Using the NetworkX package v3.4.2 (Hagberg et al. 2008) and the universal model from CarveMe (Machado et al. 2018), we built a metabolite-reaction network linking reactions through shared metabolites. Corresponding reactions for each PGFam were retrieved from the CarveMe general model, and the minimum number of metabolic steps between enzymes was determined by counting intermediate metabolites, excluding the 1,000 most common ones (e.g., water, ATP, NADH).

### Clustering and STRING-based analyses

Using the Spearman’s ⍴ values and the Phylolm estimates and *t* values from SCUE pairs, we constructed graphs using the igraph R package version 2.1.4 (Csárdi and Nepusz 2006), with these values used as edge weights. We then applied the Louvain method with a custom iterative approach to detect clusters, recursively subdividing them into smaller subclusters until no further partitioning was possible.

To link PGFams with STRING (Szklarczyk et al. 2022), we retrieved all STRING interaction gene pairs from genomes included in our dataset (Supplementary Table 1) that were also represented in the STRING database, using the STRINGdb R package version 2.16.4. For each phylum, we mapped genes to PGFams using the refseq_locus_tag identifier in BV-BRC annotations. Across all genomes, when a PGFam pair occurred multiple times, we retained only the highest combined interaction score, since we were interested in the maximum reported score. This process produced a non-redundant PGFam pair dataset of STRING interactions.

For benchmarking purposes, we used this dataset to select the highest STRING combined score *per* PGFam within each cluster, and the mean per cluster was compared between different models (Fig. 3a, Supplementary Fig. 5, Supplementary Fig. 7, and Supplementary Fig. 10).

### Analyses of duplicate genes within the same PGFam and genome

We identified cases in which multiple genes from the same genome belonged to the same PGFam, focusing on cases with exactly two genes (“duplicates”). To confirm homology, we performed BLASTp comparisons and retained only pairs with DIAMOND query and subject coverages above 90%.

Paralogs were considered within the same region (local paralogs) if there were <40 genes between them, and as remote paralogs otherwise. For remote paralogs, a distinct genetic neighborhood means that there are no common PGFams among the 20 upstream or 20 downstream genes, whereas similar genetic neighborhoods share at least two other PGFam in that region. For each duplicate, we also sampled a random gene pair from the same genome to control for genomic CUB patterns that might arise simply from co-occurence within a genome. The latter still shows weak positive correlations due to overall genome-wide CUB patterns.

### Codon and amino acid composition distances

To quantify compositional similarity between genes, we calculated codon– and amino acid–based distances using 300 randomly selected genomes from *Pseudomonadota*. We quantified distances within each genome, using only gene pairs corresponding to the PGFam pairs analyzed in our models.

Codon composition distances were computed in two steps. First, for each of the 18 amino acids with synonymous codons, we calculated the Manhattan distance as the sum of absolute differences in relative codon usage frequencies between each pair of genes. For a given amino acid, this distance equals 0 when codon usage is identical and 2 when there is no overlap in codon frequencies, regardless of the number of synonymous codons for that amino acid.

Second, the overall codon composition distance between two genes was obtained as a multidimensional Euclidean distance, calculated as the square root of the sum of squared amino–acid specific distances. The resulting values range from 0 (identical codon composition) to √72 ≃ 8.48, corresponding to no overlap in codon usage.

For amino acid composition, relative amino acid frequencies were computed for each gene, and Euclidean distances were calculated between all gene pairs, yielding values ranging from 0 (identical amino acid composition) to 1 (no shared amino acid usage).

## Figures and code availability

All figures were generated in R (https://www.r-project.org/) using ggplot2 (v3.5.2) and, when necessary, edited in InkScape. Analyses were performed with custom R scripts, available at Zenodo (López at al. 2025).

## Author contributions

Conceptualization: JLL, BED Methodology: JLL, MDB, AL Investigation: JLL Visualization: JLL

Funding acquisition: BED Project administration: BED Resources: BED Supervision: BED, JLL Writing - original draft: JLL

Writing - review & editing: JLL, BED Software: JLL, AL

Data curation and validation: SPD, JLL, MS, MDB, AL, EP Formal analysis: JLL, BED, MS, AL

## Data and materials availability

All data and code generated and/or analyzed in this study are deposited at Zenodo (López at al. 2025)

