## Supplementary Fig for "Widespread synchronization of codon usage in functionally related genes"

### Supplementary Material

**The PDF file includes:**

Supplementary Notes

Supplementary Figs. 1 to 17

### Supplementary Notes

##### *Supplementary Note 1 – GCB and GCB_mod are the CUB indices that best reflect expression*

As codon usage reflects expression (Hanson and Coller 2018), we benchmarked seven CUB indices against gene expression levels across a range of experimental conditions. Expression data were obtained from the Precise 1K dataset from *Escherichia coli* K-12 MG1655 (Sastry et al. 2019). Among the tested indices, the gene codon bias (GCB) index showed the strongest and most consistent correlation with gene expression across genomes compared with other commonly used indices (Supplementary Fig. 1 and Supplementary Fig. 2).

The selected GCB index initially uses the set of ribosomal protein genes as a seed reference set, estimates GCB values for all genes in the genome, and then iteratively searches for a new optimal reference set comprising the genes with the highest GCB values, until reaching a local maximum (Merkl 2003). For our first analyses shown in Fig. 1, we retained only those genomes in which at least 50% of ribosomal protein genes were included in the final optimized reference set. This filtering was necessary because, in most genomes, the iterative optimization converged to an alternative local maximum that did not include the ribosomal genes. Thus, the genomes used in the initial analyses (Figs. 1–2) were those meeting this criterion and are listed as “Main phyla - small set” in Table S1. Comparable results were obtained using a modified version of the index (GCB_mod; Supplementary Fig. 1 and Supplementary Fig. 2), which enabled the inclusion of additional reference species and increased statistical power in our SCUE analyses. GCB_mod simply calculates the initial GCB index without any iteration. We therefore used GCB_mod for the rest of the study (Figs. 3–4 and Supplementary Figs. 4 onwards), as it allowed the inclusion of a larger and more phylogenetically diverse dataset, yielding more clusters and incorporating additional phyla. Interestingly, GCB and GCB mod performed better than the commonly used CAI index (Sharp and Li 1987), suggesting that this index may be more appropriate for CU studies.

***Supplementary Note 2 – Most duplicated PGFams conserve similar CUB within the same genome***

To examine how homologs within the same genome adapt their CUB, we analyzed PGFams containing multiple gene copies per genome. We found that 75-88% of PGFams included at least one genome with more than one copy. Although some PGFams showed a high number of gene copies per genome, such as those encoding transposases and other mobile genetic elements, ABC transporters, two-component system proteins, transcriptional regulators, motility and chemotaxis-related proteins, Type VI secretion system components, and RND efflux system proteins, most multiple copy PGFams were represented genes with by two-copies (Supplementary Fig. 17).

We detected between 12,184 and 44,381 cases of duplicated genes across 267 to 1,122 genomes and 568 to 1,896 PGFams among the main five phyla, with averages ranging from 1.1 to 3.8 duplicates per genome. To focus on true isofunctional homologs, we retained only those gene pairs with > 90% query and subject coverage in DIAMOND blastp searches, which represented 7.1-14.1% of all detected duplicates. When comparing GCB values among duplicates located in non-overlapping genomic regions, we observed a positive and significant correlation across the five phyla (Supplementary Fig. 3). To rule out potential biases arising from genome-wide CUB patterns shared by duplicated genes, we repeated the same analyses using randomized sets of gene pairs from the same genomes. These random pairs also showed positive but significantly lower correlations than the duplicates. Across all five phyla, duplicated genes had significantly higher $\rho$ values compared to random pairs (Fisher's Z-transformation test, $p$ value < 0.05, Supplementary Table 2), indicating that functional constraints, rather than genomic context alone, primarily drive the conservation of CUB optimization among duplicates. We also found that correlations were stronger for duplicates located within the same genomic region, or for those positioned distantly but sharing neighboring PGFams, likely reflecting functional conservation mediated by homologous recombination that preserved ancestral CUB patterns. Importantly, even after removing these cases and retaining only duplicates without shared neighboring PGFams, correlations remained significantly positive (Supplementary Fig. 3). Altogether, these results indicate that duplicated genes generally preserve their CUB optimization patterns, suggesting that codon usage adaptation is maintained following gene duplication due to functional constraints and not only due to shared ancestry.

#### Figures

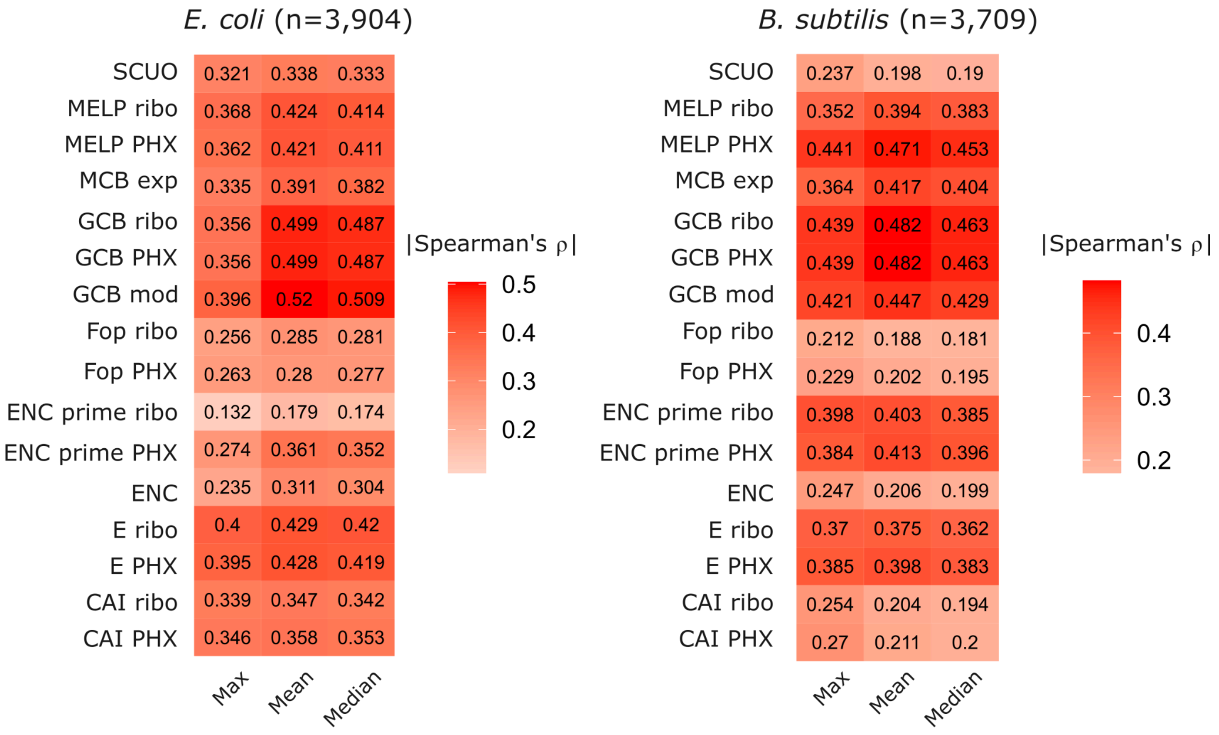


**Supplementary Fig. 1.**

**Spearman correlations between CUB indices and gene expression levels.** To benchmark the CUB indexes (rows), transcriptional data from the Precise 1K dataset from *Escherichia coli* K-12 MG1655 (Sastry et al. 2019) and MODULOME data from *Bacillus subtilis* 168 (Sastry et al. 2024) were summarized using different statistical measures, and absolute Spearman correlations between CUB indexes and gene expression were calculated. CUB indices were computed with the coRdon package (Elek 2019) using two sets of putatively highly expressed reference genes: ribosomal proteins (“ribo”) and a manually curated set (“PHX”) comprising genes with high protein expression in the PaxDB database (Wang et al. 2015) in at least two of the three species. Maximum, mean, and median expression values were calculated from TPM-normalized expression values across a wide range of conditions. For subsequent analyses, we selected GCB (Merkl 2003) and GCBmod, both calculated using ribosomal proteins as the reference set, as this approach showed the strongest correlations and was simpler to implement than the PHX-based method.

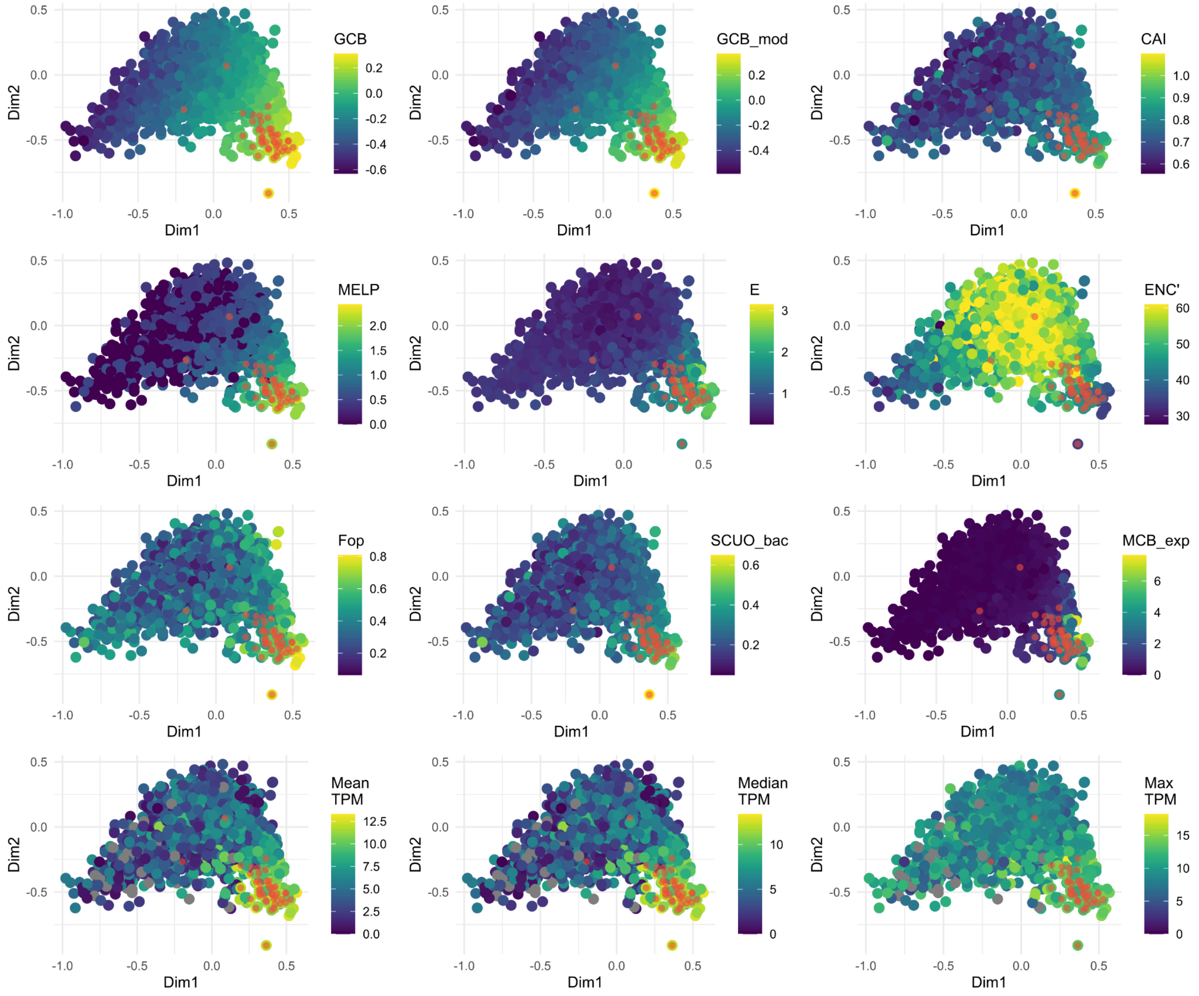


**Supplementary Fig. 2.**

**Factorial correspondence analysis of the *Escherichia coli* K-12 MG1655 genome.** Different CUB indices (GCB, GCB_mod, CAI, MELP, E, ENC', Fop, SCUO, and MCB) are shown in distinct colors to illustrate how each index represents gene-level CUB. Ribosomal genes (red dots), were used as the reference set of highly expressed genes. Maximum, mean, and median expression values were calculated from TPM-normalized expression data across a wide range of conditions obtained from the Precise 1K dataset from *Escherichia coli* K-12 MG1655 (Sastry et al. 2019). Among the tested indices, GCB decreases most gradually from ribosomal genes and shows the strongest correlation with expression data (Supplementary Fig. 1). All indices were calculated using ribosomal genes as the reference. Gray dots denote genes lacking expression data.

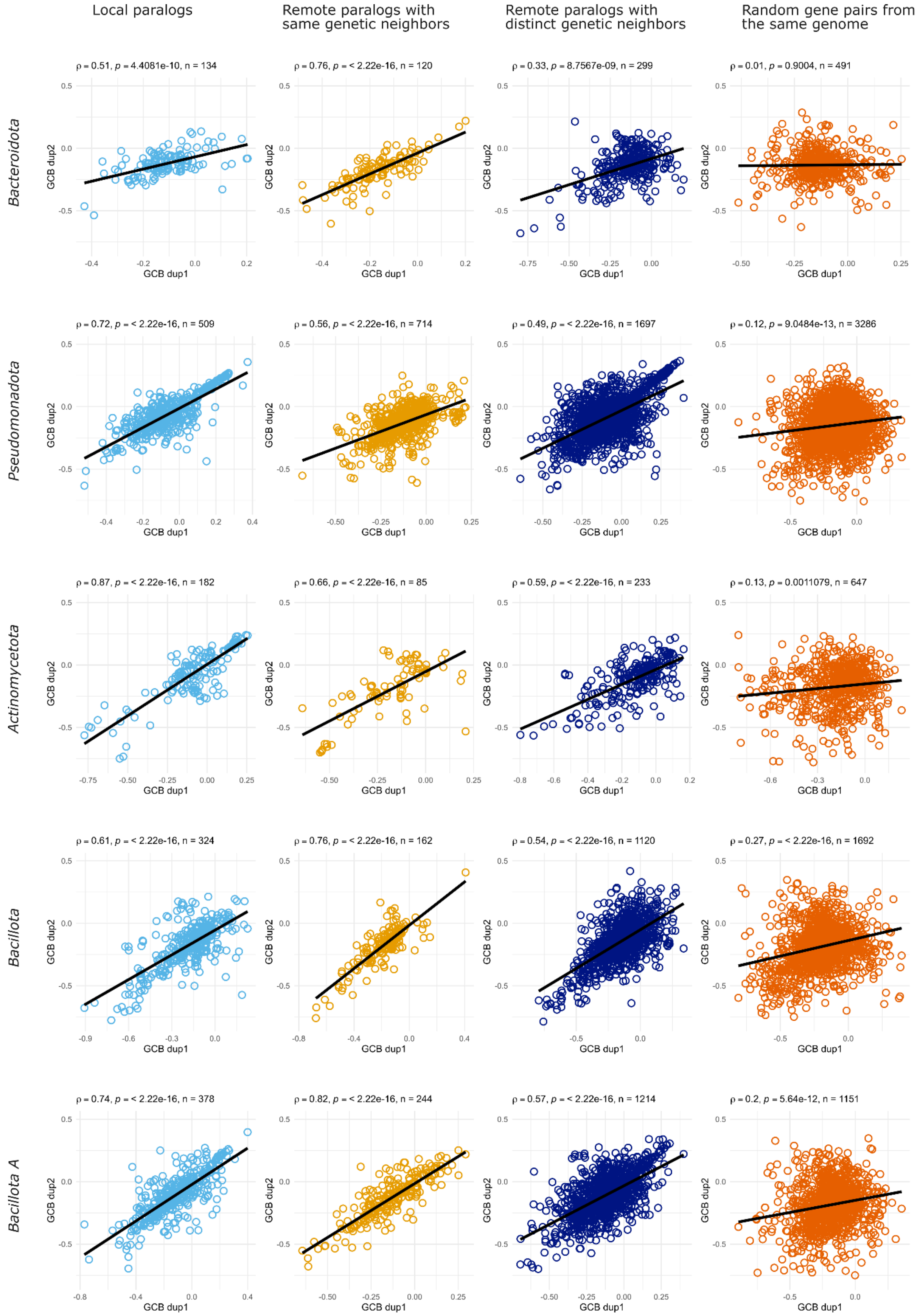


**Supplementary Fig. 3.**

**Correlation between duplicate genes from the same PGFam and genome in main phyla.** Both genomically co-localized paralogs, and remote paralogs with similar genomic neighbourhoods present the highest correlation values, while distant duplicates without evident synteny present a lower but still significant positive correlation.

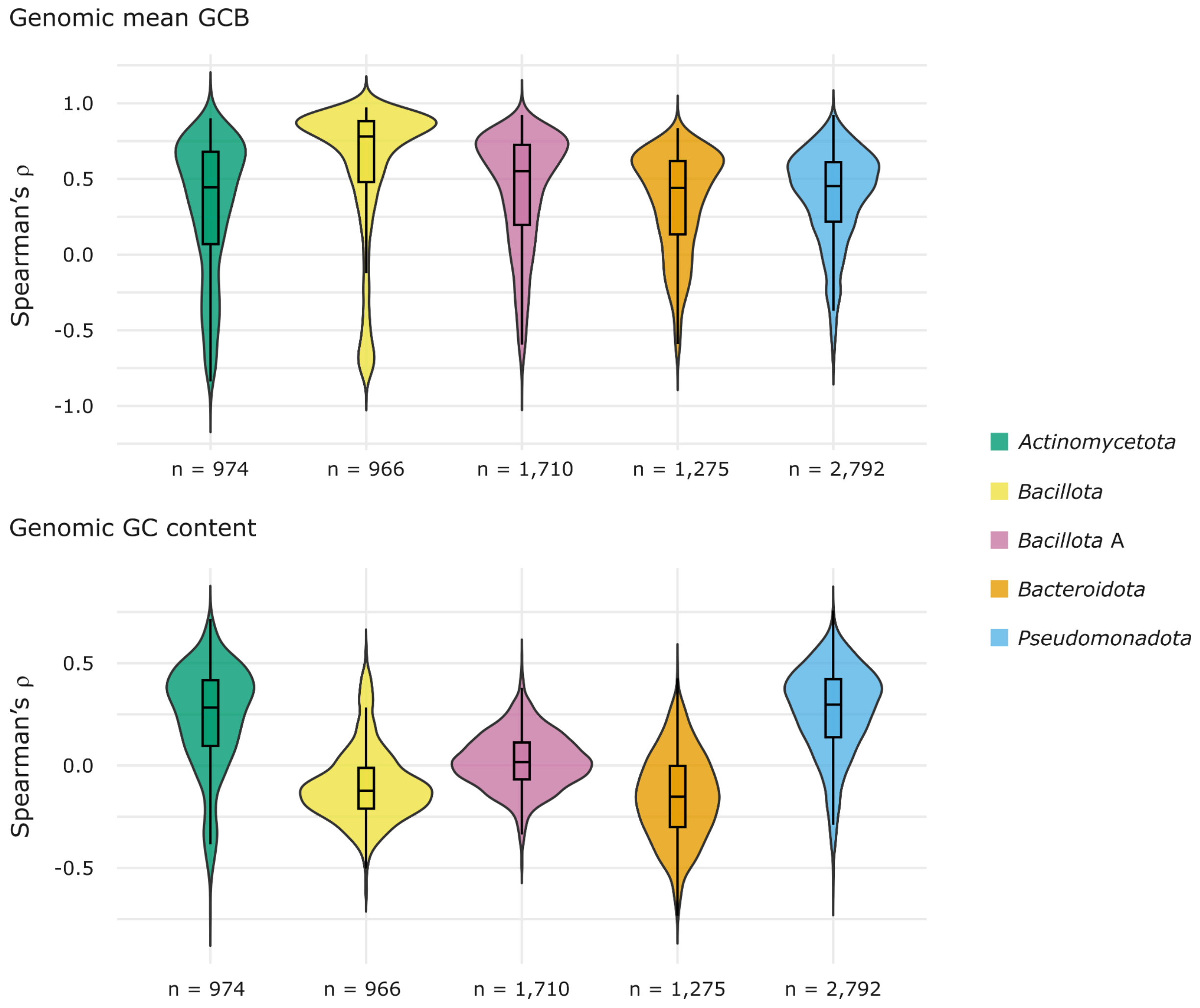


**Supplementary Fig. 4.**

**Genomic mean GCB and GC content in genomes are highly correlated with GCB and are factors that influence correlation values.** Spearman correlations were calculated between single-copy PGFam GCB values and the corresponding genomic mean CUB or GC content and their $\rho$ values are shown for each phyla (Supplementary Table 2). Only PGFams present in more than 100 genomes from the small genome set (Supplementary Table 1) were used for the correlations, whose number of correlations (n) is indicated for each phylum.

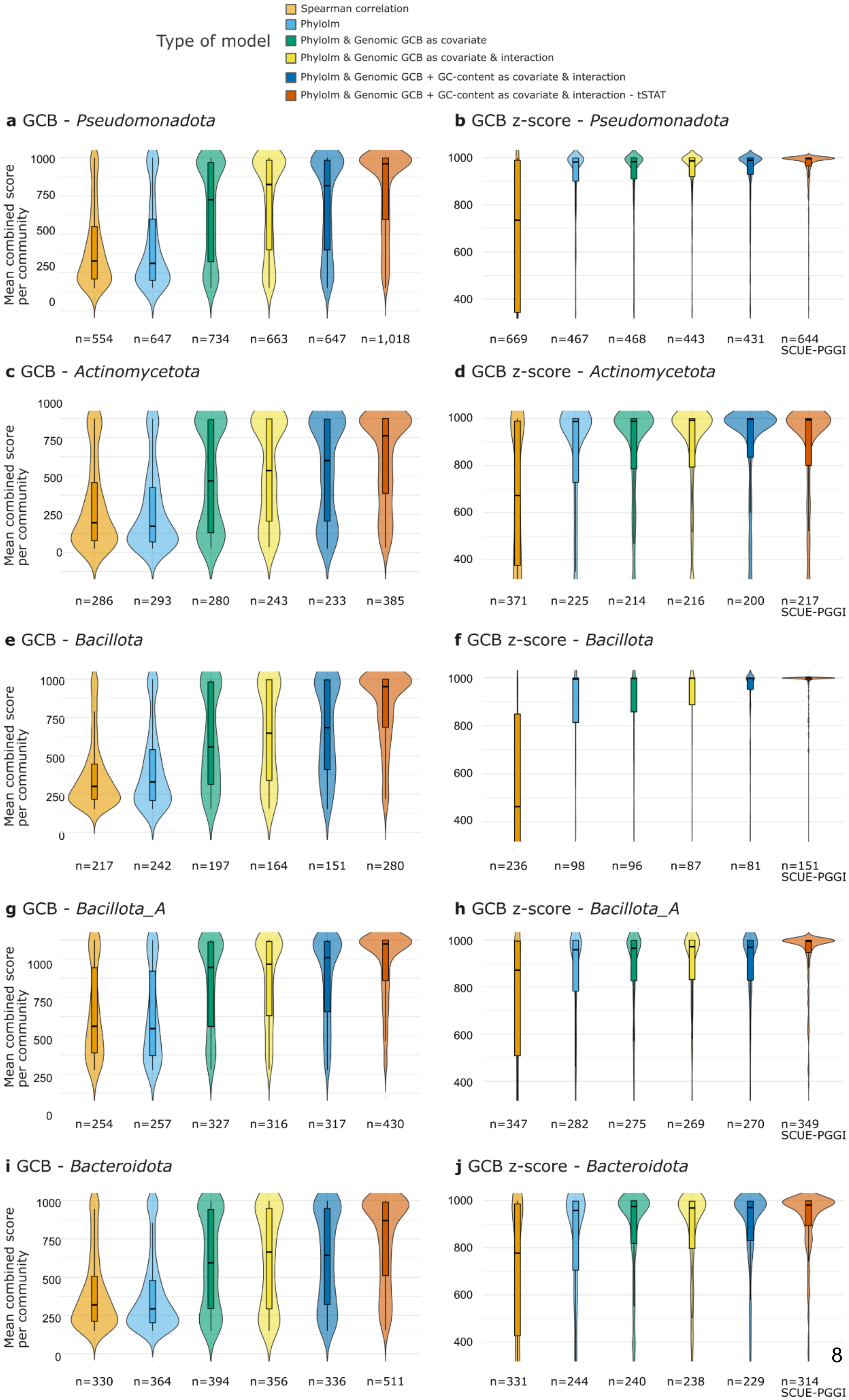
Type of model
Spearman correlation
Phylolm
Phylolm & Genomic GCB as covariate
Phylolm & Genomic GCB as covariate & interaction
Phylolm & Genomic GCB + GC-content as covariate & interaction
Phylolm & Genomic GCB + GC-content as covariate & interaction - tSTAT
a GCB - Pseudomonadota
b GCB z-score - Pseudomonadota
Mean combined score per community
n=554 n=647 n=734 n=663 n=647 n=1,018
n=669 n=467 n=468 n=443 n=431 n=644
SCUE-PGGI
c GCB - Actinomycetota
d GCB z-score - Actinomycetota
n=286 n=293 n=280 n=243 n=233 n=385
n=371 n=225 n=214 n=216 n=200 n=217
e GCB - Bacillota
f GCB z-score - Bacillota
n=217 n=242 n=197 n=164 n=151 n=280
n=236 n=98 n=96 n=87 n=81 n=151
g GCB - Bacillota_A
h GCB z-score - Bacillota_A
n=254 n=257 n=327 n=316 n=317 n=430
n=347 n=282 n=275 n=269 n=270 n=349
i GCB - Bacteroidota
j GCB z-score - Bacteroidota
n=330 n=364 n=394 n=356 n=336 n=511
n=331 n=244 n=240 n=238 n=229 n=314

**Supplementary Fig. 5.**

**Clusters obtained with iterative Louvain using different correlation and phylolm models.** GCB (**a,c,e,g,i**) or GCB z-score (**b,d,f,h,j**) values for each pair of PGFams across genomes were used as input for Spearman correlation or phylolm regression models. We used phylolm models with a GTDB based tree as a phylogenetic control, and Genomic GCB of each genome as a covariate, with and without an interaction term to our independent variable. Spearman's $\rho$ values or phylolm estimates or *t* values from the phylolm models were used as weights to generate networks using the igraph R package. An iterative Louvain clustering approach was subsequently applied and resulting clusters were mapped to STRING interactions in these genomes. We obtained more biologically meaningful clusters (higher mean STRING combined scores) using GCB z-scores, and including the genomic GCB and GC content as covariate and interaction. N represents the number of clusters obtained with at > 2 members having STRING interaction scores. SCUE-PGGI is the model using *t* values from phylolm instead as estimates, including covariates and interactions.

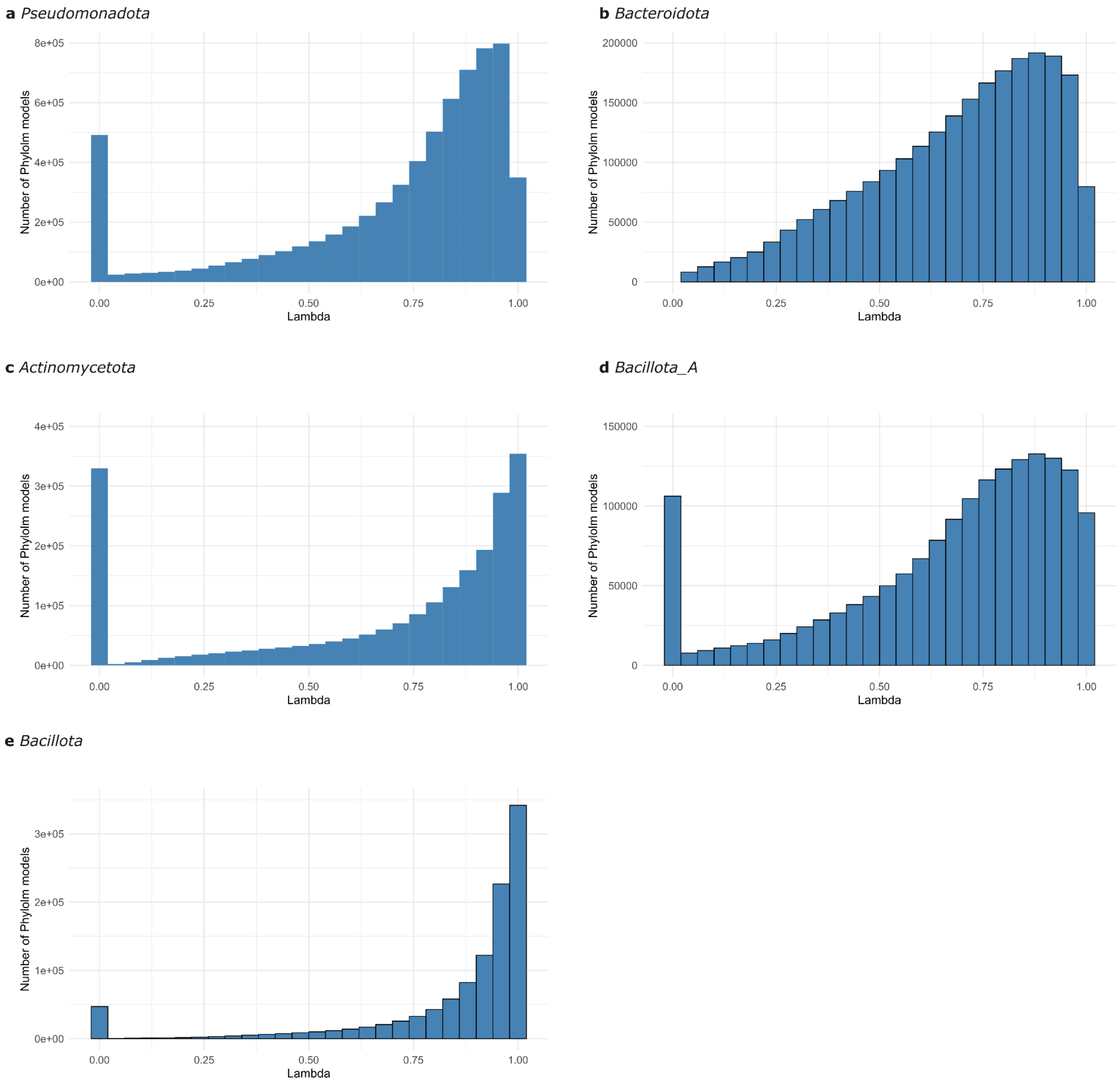


**Supplementary Fig. 6.**

**Histogram of lambda values in SCUE-P models.** Pagel's λ values were extracted from the output of each SCUE-P model and are displayed as histograms. Values approaching 1 indicate a strong phylogenetic signal in the models.

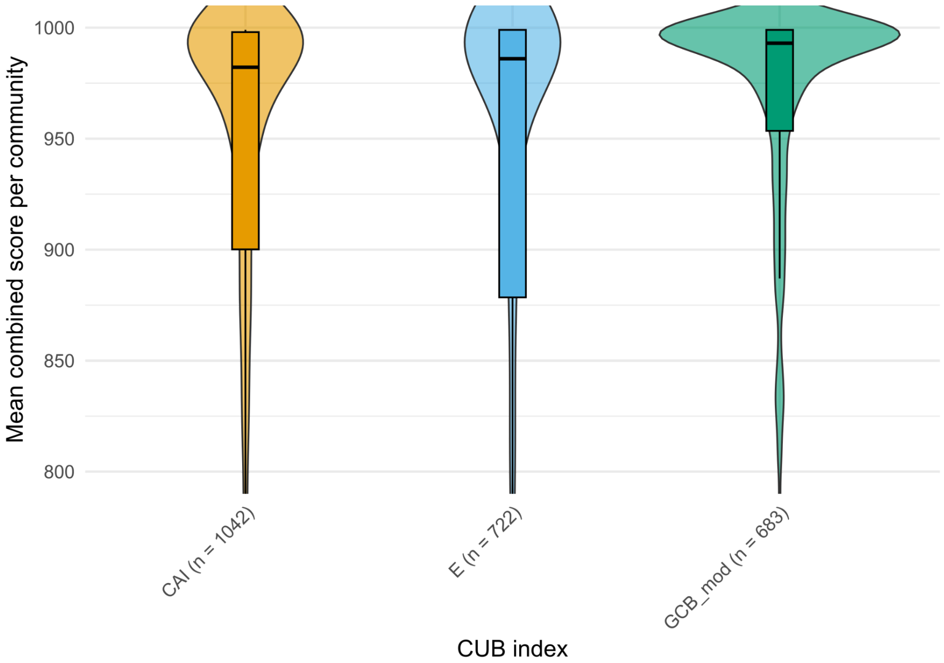


**Supplementary Fig. 7. Clusters obtained with iterative Louvain using SCUE-P models with different CUB indices.** CAI z-score (**a**), E z-score (**b**), or GCB z-score values for each pair of PGFams across genomes were used as input for phylolm regression models (SCUE-P). We used phylolm models with a GTDB based tree as a phylogenetic control. Phylolm *t* values from the phylolm models were used as weights to generate networks using the igraph R package. An iterative Louvain clustering approach was subsequently applied and resulting clusters were mapped to STRING interactions in these genomes. For each PGFam inside a cluster, we only retained the highest STRING interaction. We obtained more biologically meaningful clusters (higher mean STRING combined scores) using GCB z-scores, instead of CAI z-scores or E z-scores, although more clusters were obtained with the latter. N represents the number of clusters obtained with at > 2 members having STRING interaction scores.

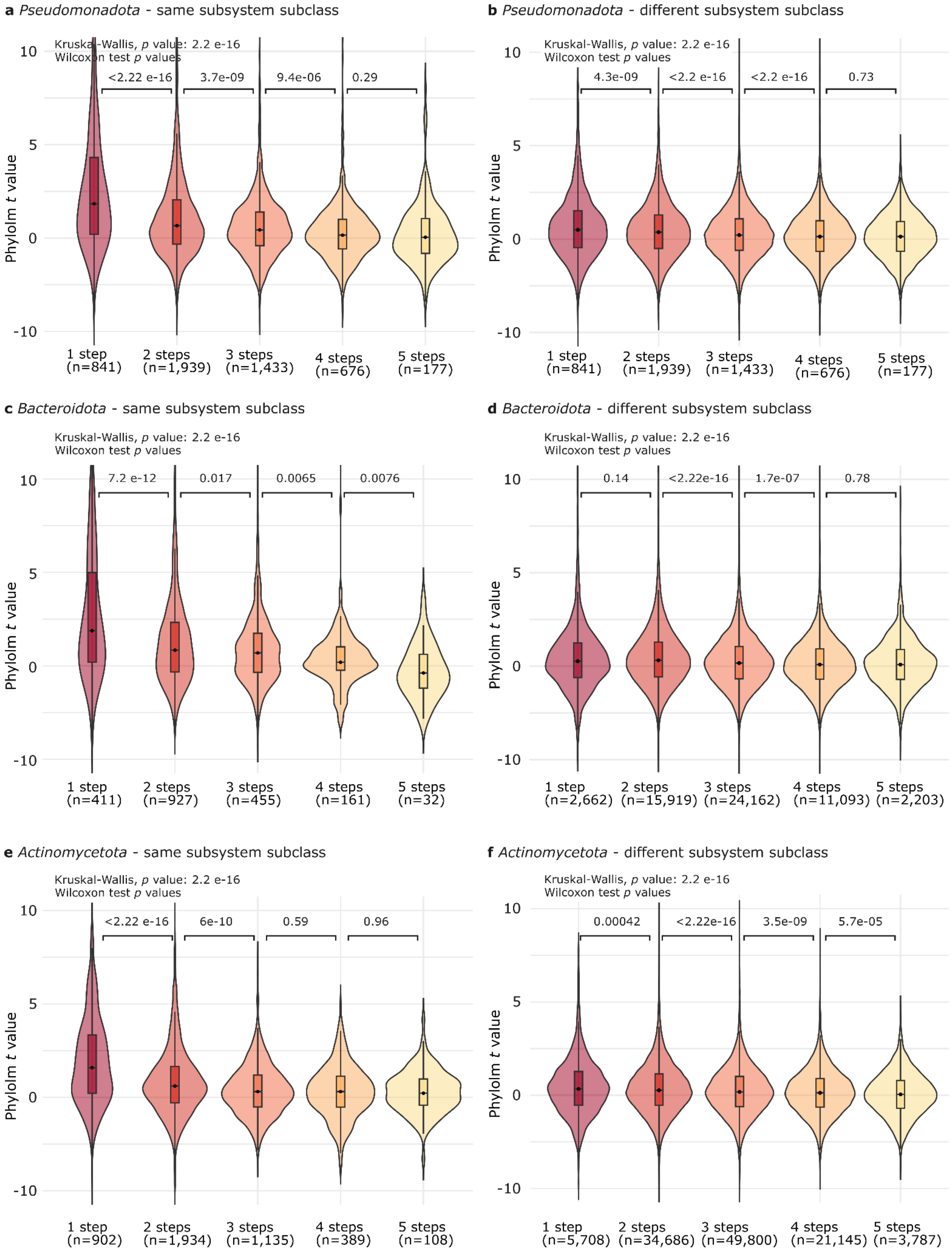
a Pseudomonadota - same subsystem subclass
Kruskal-Wallis, p value: 2.2 e-16
Wilcoxon test p values
<2.22 e-16
3.7e-09
9.4e-06
0.29
Phylolm t value
10
5
0
-10
1 step (n=841)
2 steps (n=1,939)
3 steps (n=1,433)
4 steps (n=676)
5 steps (n=177)
b Pseudomonadota - different subsystem subclass
Kruskal-Wallis, p value: 2.2 e-16
Wilcoxon test p values
4.3e-09
<2.2 e-16
<2.2 e-16
0.73
Phylolm t value
1 step (n=841)
2 steps (n=1,939)
3 steps (n=1,433)
4 steps (n=676)
5 steps (n=177)
c Bacteroidota - same subsystem subclass
Kruskal-Wallis, p value: 2.2 e-16
Wilcoxon test p values
7.2 e-12
0.017
0.0065
0.0076
Phylolm t value
1 step (n=411)
2 steps (n=927)
3 steps (n=455)
4 steps (n=161)
5 steps (n=32)
d Bacteroidota - different subsystem subclass
Kruskal-Wallis, p value: 2.2 e-16
Wilcoxon test p values
0.14
<2.22e-16
1.7e-07
0.78
Phylolm t value
1 step (n=2,662)
2 steps (n=15,919)
3 steps (n=24,162)
4 steps (n=11,093)
5 steps (n=2,203)
e Actinomycetota - same subsystem subclass
Kruskal-Wallis, p value: 2.2 e-16
Wilcoxon test p values
<2.22 e-16
6e-10
0.59
0.96
Phylolm t value
1 step (n=902)
2 steps (n=1,934)
3 steps (n=1,135)
4 steps (n=389)
5 steps (n=108)
f Actinomycetota - different subsystem subclass
Kruskal-Wallis, p value: 2.2 e-16
Wilcoxon test p values
0.00042
<2.22e-16
3.5e-09
5.7e-05
Phylolm t value
1 step (n=5,708)
2 steps (n=34,686)
3 steps (n=49,800)
4 steps (n=21,145)
5 steps (n=3,787)

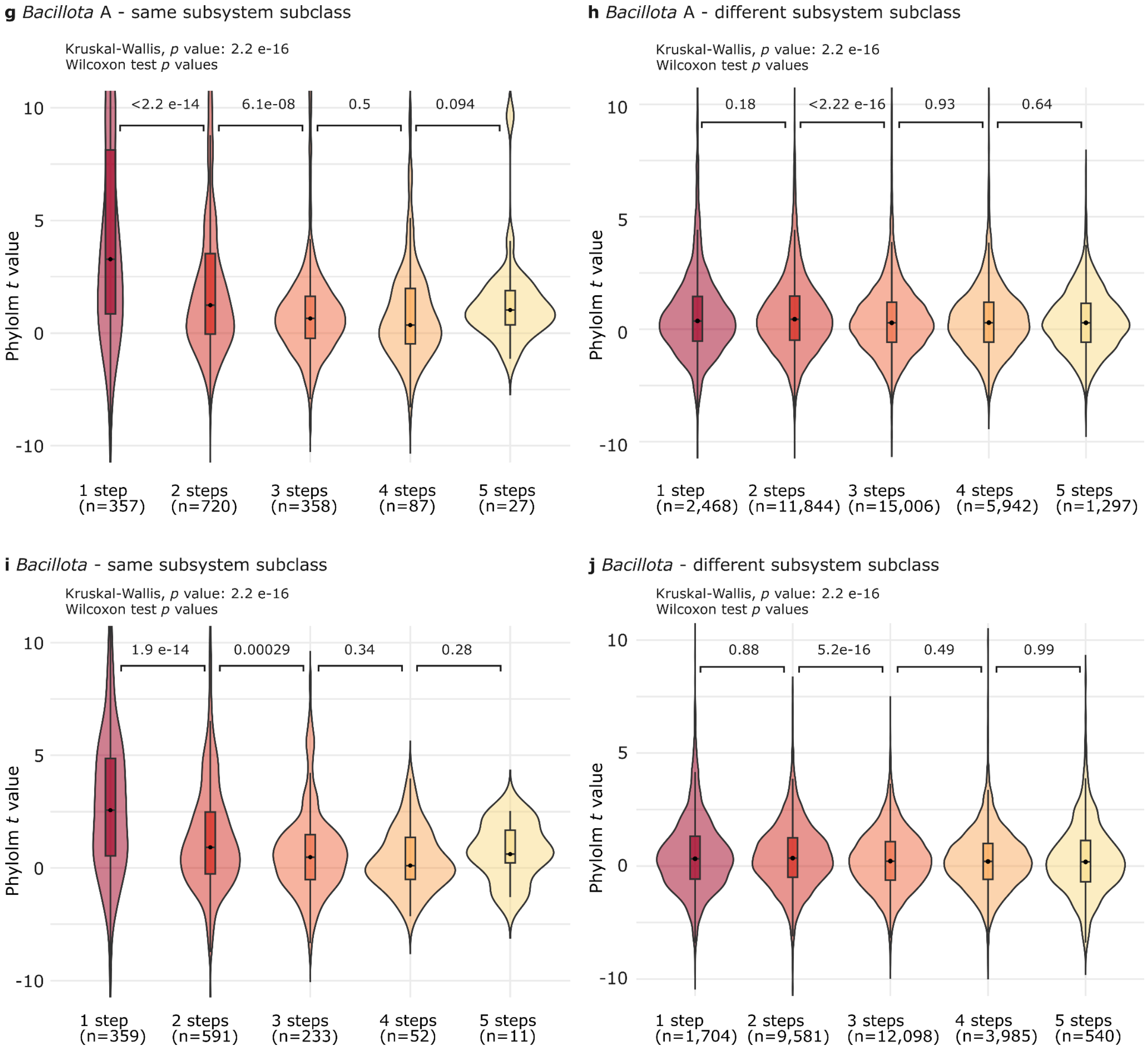


**Supplementary Fig. 8.**

**SCUE is higher in closer enzymes in metabolism, especially when classified to the same subsystem.**

SCUE-PGGI derived phylolm *t* values increase when genes from the same (**a,c,e,g,i**) and different (**b,d,f,h,j**) subsystem subclass are closer in metabolic pathways (enzymes 1 step apart in metabolic networks) compared to more distant genes (5 steps apart).

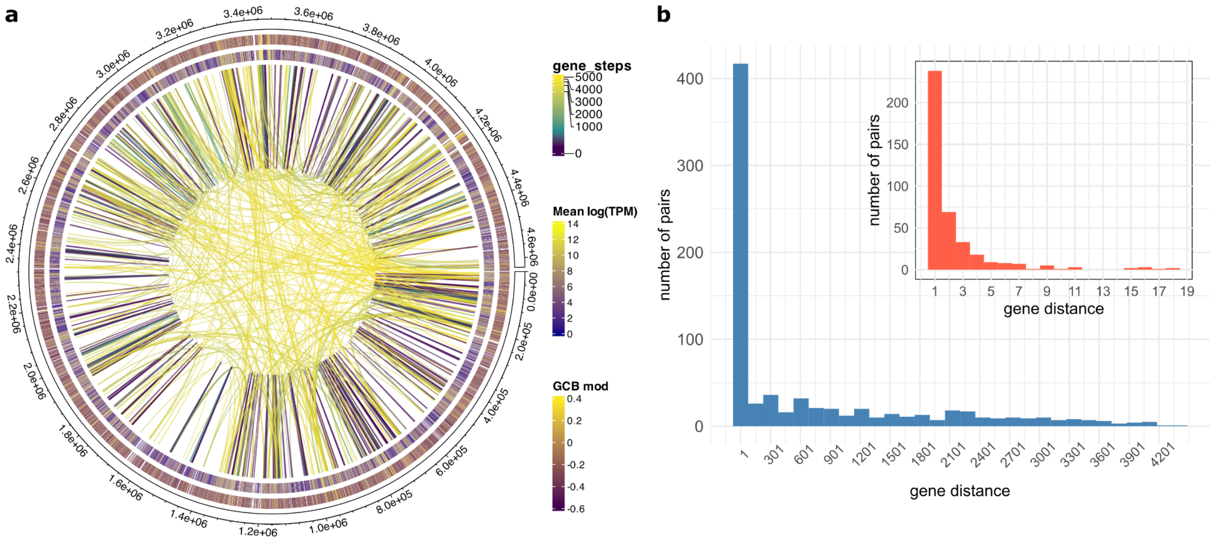


**Supplementary Fig. 9.**

**Significant pairs link both close and distant genes. a** Significant SCUE pairs (FDR <0.0001) in *Pseudomonadota* were plotted using the circlize R package (v. 0.4.17) to display connections colored by the number of genes separating the pair (gene steps, inner links) on the *Escherichia coli* K-12 MG1655 genome. Colored strip rings represent the mean log(TPM) expression values (inner ring) from the Precise 1K dataset, and the GCB mod values of genes (outer ring). **b** The distribution of the number of pairs with a gene distance within each bin is represented. We represented gene distance as the number of gene steps to reach the second gene. Each PGFam was represented with only the pair with highest *t* value.

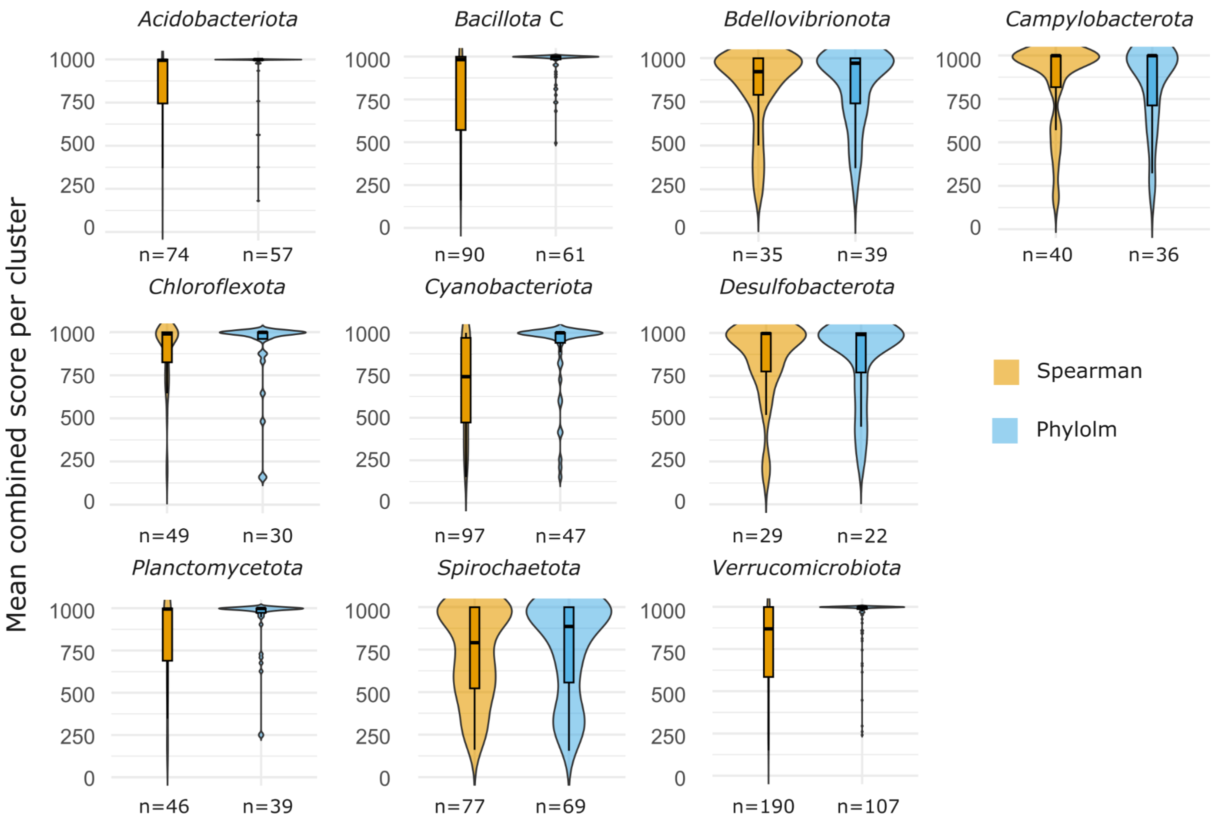


**Supplementary Fig. 10.**

**Clusters obtained with iterative Louvain across an extended set of phyla.** GCB z-score values for each pair of PGFam across genomes were used as input for Spearman correlation, or SCUE-P models. We used a GTDB based tree as a phylogenetic control. Spearman's $\rho$ values or SCUE-P *t* values were used as weights to generate networks using the igraph R package. An iterative Louvain clustering approach was subsequently applied and resulting clusters were mapped to STRING interactions in these genomes. N represents the number of clusters obtained with members having known interactions within clusters (n of members > 2).

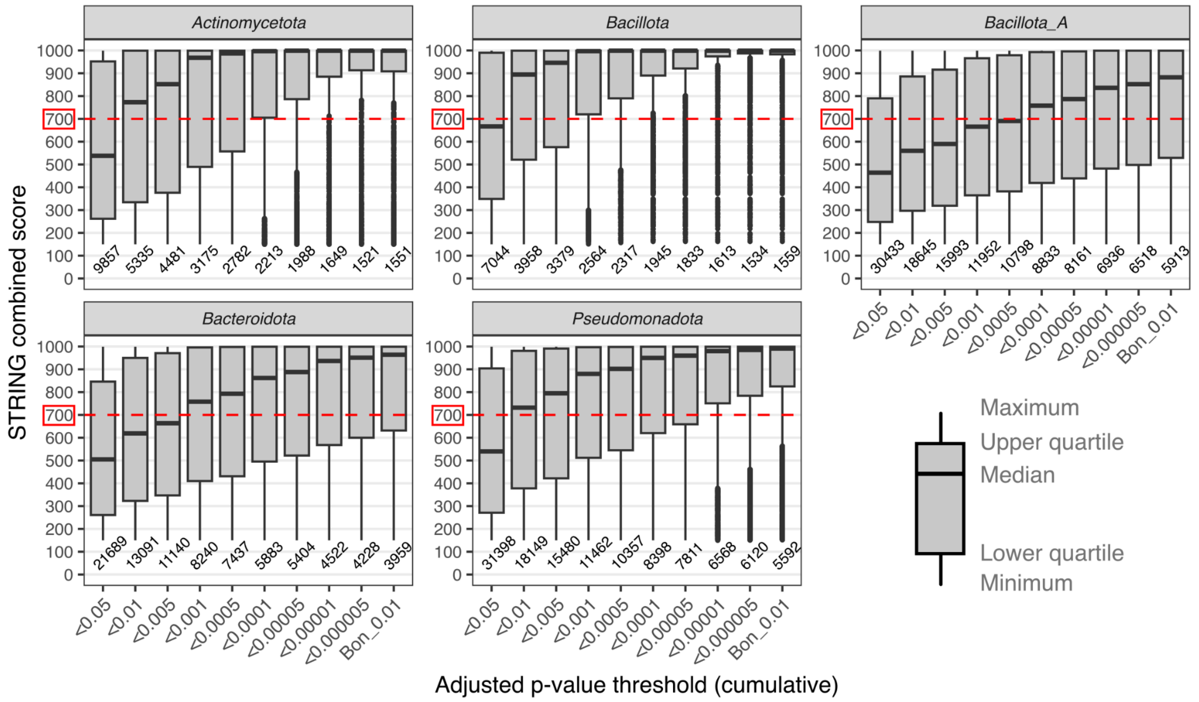


**Supplementary Fig. 11.**

**Comparison of *p* adjusted thresholds in SCUE-PGGI *t* values.** Different Benjamini-Hochberg (BH) p adjusted cutoffs were applied to SCUE pairs which were subsequently mapped to their STRING combined scores. X axis denote the BH cutoff values and the Bonferroni stringent cutoff (Bon_0.01 for Bonferroni p adjusted <0.01). Each phylum results in a different optimal cutoff where its median combined score reaches a value of ca. 700 (i.e., *Actinomycetota* cutoff < 0.01, Supplementary Table 4). The box represents the interquartile range (IQR: 25th–75th percentile), the horizontal line inside the box is the median, and the whiskers extend to the minimum and maximum non-outlier values.

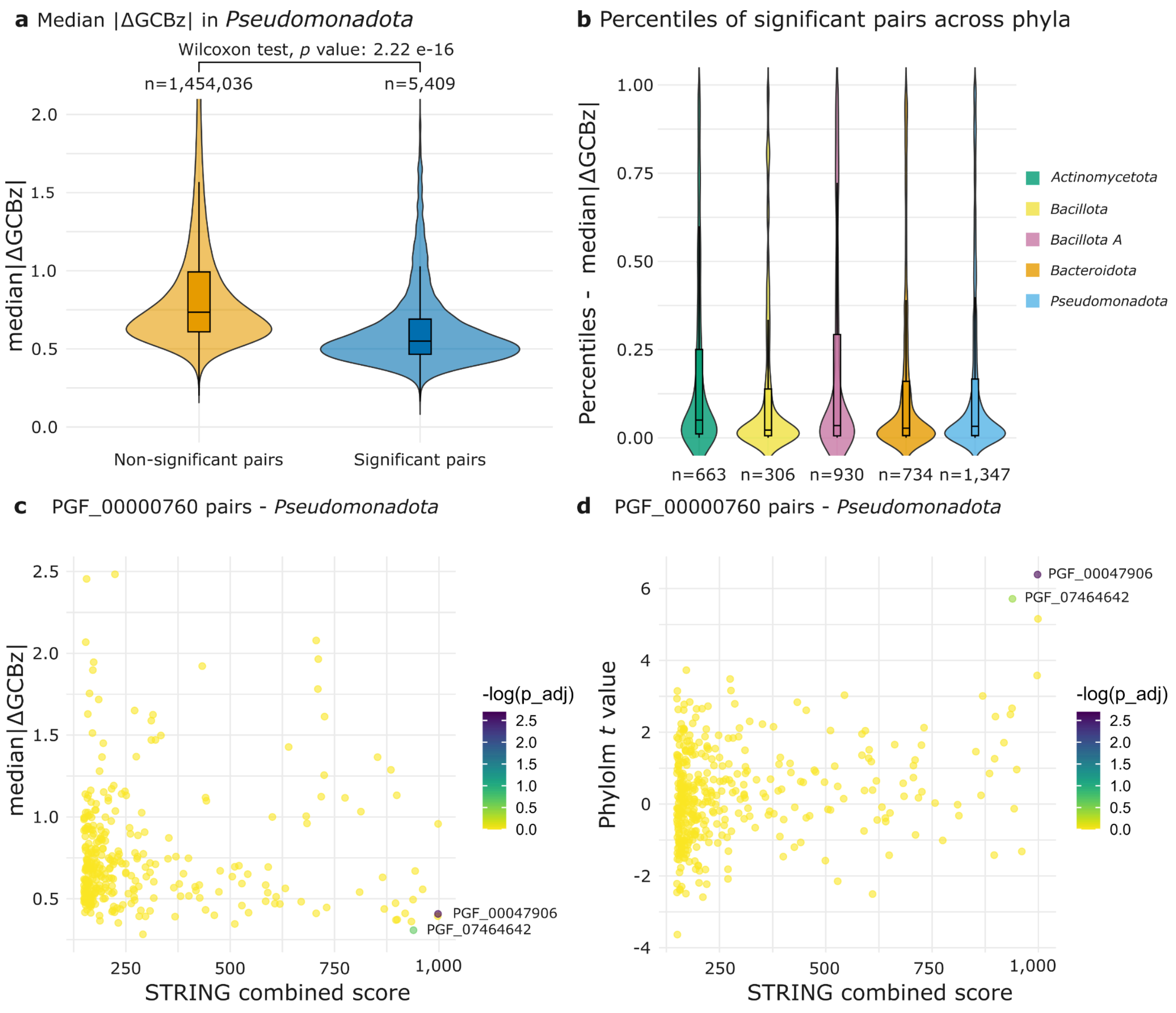


**Supplementary Fig. 12.**

**Low median differences in GCB z-scores (i.e., |ΔGCBz|) serves as an additional indicator of SCUE. a** Median |ΔGCBz| values are significantly lower in significantly SCUE pairs, indicating that they have more similar CUB across genomes than pairs that do not co-evolve. **b** For each PGFam containing significantly SCUE pairs, the most significant pair typically ranks among the lowest percentiles of median |ΔGCBz| compared to all other pairs from the same PGFam. The distribution of these percentiles across the five main phyla is shown. n indicates the number of significant pairs analyzed. **c**,**d** Example of SCUE pairs associated with PGF_00000760 (α-glucoside ABC transporter, permease protein AglF). The two top-scoring pairs are PGF_00047906 (α-glucoside ABC transporter, permease protein AglG) and PGF_07464642 (maltodextrin glucosidase, EC 3.2.1.20), both showing low median |ΔGCBz| **c** and high phylolm *t* values **d** compared to all other pairs. While PGF_07464642 is non-significant under Bonferroni correction (*p* adjusted <0.01), it becomes significant when using a more relaxed FDR correction (*p* adjusted <0.05).

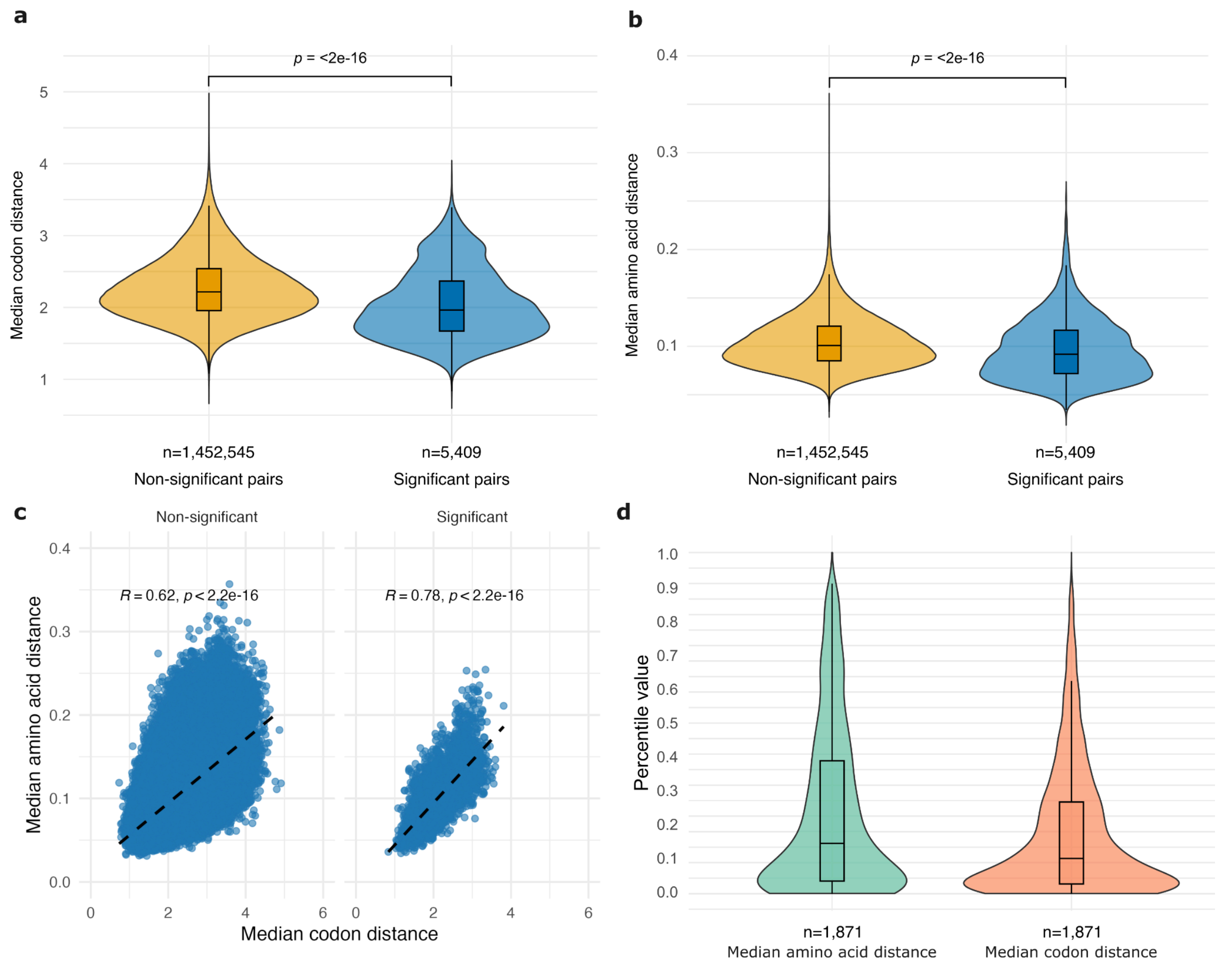


**Supplementary Fig. 13.**

**Codon and amino acid distances in SCUE pairs.** Median **a** codon and **b** amino acid distances are significantly lower in significant (SCUE) compared to non-significant gene pairs. Distances were calculated using gene pairs corresponding to PGFams included in our models that presented at least one significant pair (n=1,826), from 300 randomly selected genomes affiliated with *Pseudomonadota*. Significant pairs were considered those with *p* adjusted (Bonferroni) values < 0.01. The Wilcoxon test was used to compare the distances between significant and non-significant gene pairs. N represents the number of paired distances in each group, derived from the 300 genomes. **c** Spearman correlation plots between median codon distances and median amino acid distances for significant and non-significant pairs. Significant pairs show a higher correlation (R=0.77 *vs* 0.6) between codon and amino acid distances. **d** For each PGFam containing significant SCUE pairs, the most significant pair typically ranks among the lowest percentiles of codon and amino acid distances compared to all other pairs from the same PGFam. The distribution of these percentiles within genomes affiliated with *Pseudomonadota* are shown. n indicates the number of significant pairs analyzed.

de multiple PGFams with > 1 representative.

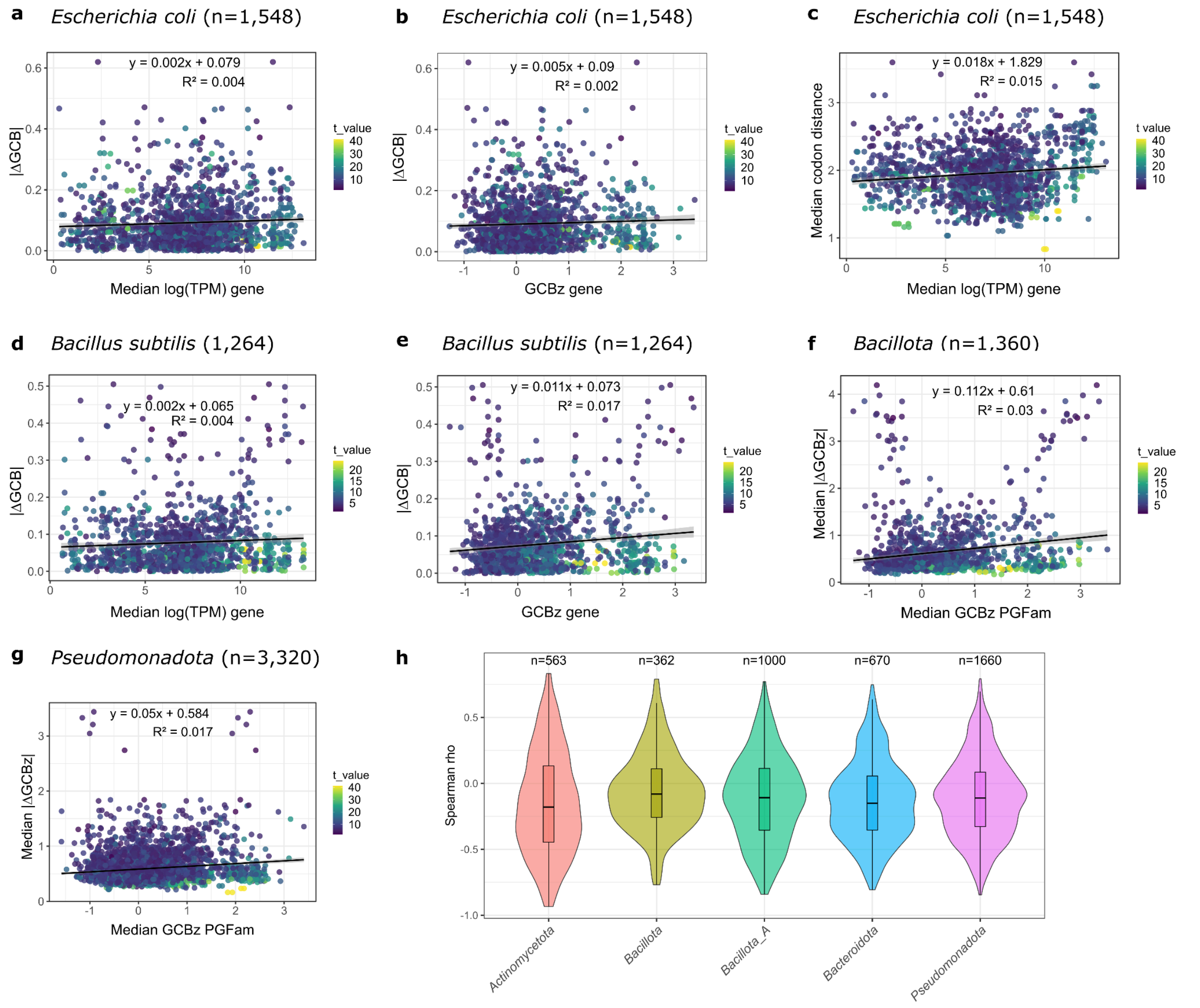


**Supplementary Fig. 14**

**Codon usage distance is similar both in lowly and highly expressed significant gene pairs.** |ΔGCB| values *vs*. expression (as median logTPM values across all conditions from the **a** Precise 1K dataset in *E. coli* (Sastry et al. 2019) or the **d** MODULOME dataset in *B. subtilis* (Sastry et al. 2024), and GCBz values are shown for genes in positive significant pairs (*t* value > 0, FDR < 0.0001) in *E. coli* and *B. subtilis* (**b**, **e**, respectively). For each PGFam, significant pairs were filtered to show only the pair with the highest *t* value across genomes in *Pseudomonadota* and *Bacillota*, respectively. The representative gene of each PGFam was selected in the *E. coli* K-12 MG1655 and *B. subtilis* subsp. *subtilis* str. 168 genomes. Each dot represents the expression or GCBz value of each gene in pairs (x-axis) and the median |ΔGCB| value of each pair (y-axis). Each pair is represented twice to include both genes in the pair. **c** Median codon distance *vs*. expression. **f**, **g** Median |ΔGCBz| values and *vs.* Median GCB *z* values of each gene within selected significant pairs across all genomes in *Bacillota* and *Pseudomonadota*, respectively, illustrate the same observation can be extended to all significant pairs in the phyla. **h** Significant pairs were individually selected, genes representing their PGFams were obtained for each genome where the pair was present, and for each pair a Spearman correlation was calculated between the GCBz values of genes and the |ΔGCBz| of their corresponding pairs across genomes. Then, *rho* values of each of these correlations were represented in a violin plot. Numbers indicate the number of individual correlations. Only one correlation per pair was calculated with one randomly selected gene per pair. As the distribution of correlations is lower than 0, this evidence shows that in significant pairs the |ΔGCBz| values are in general higher in lowly expressed compared to highly expressed genes.

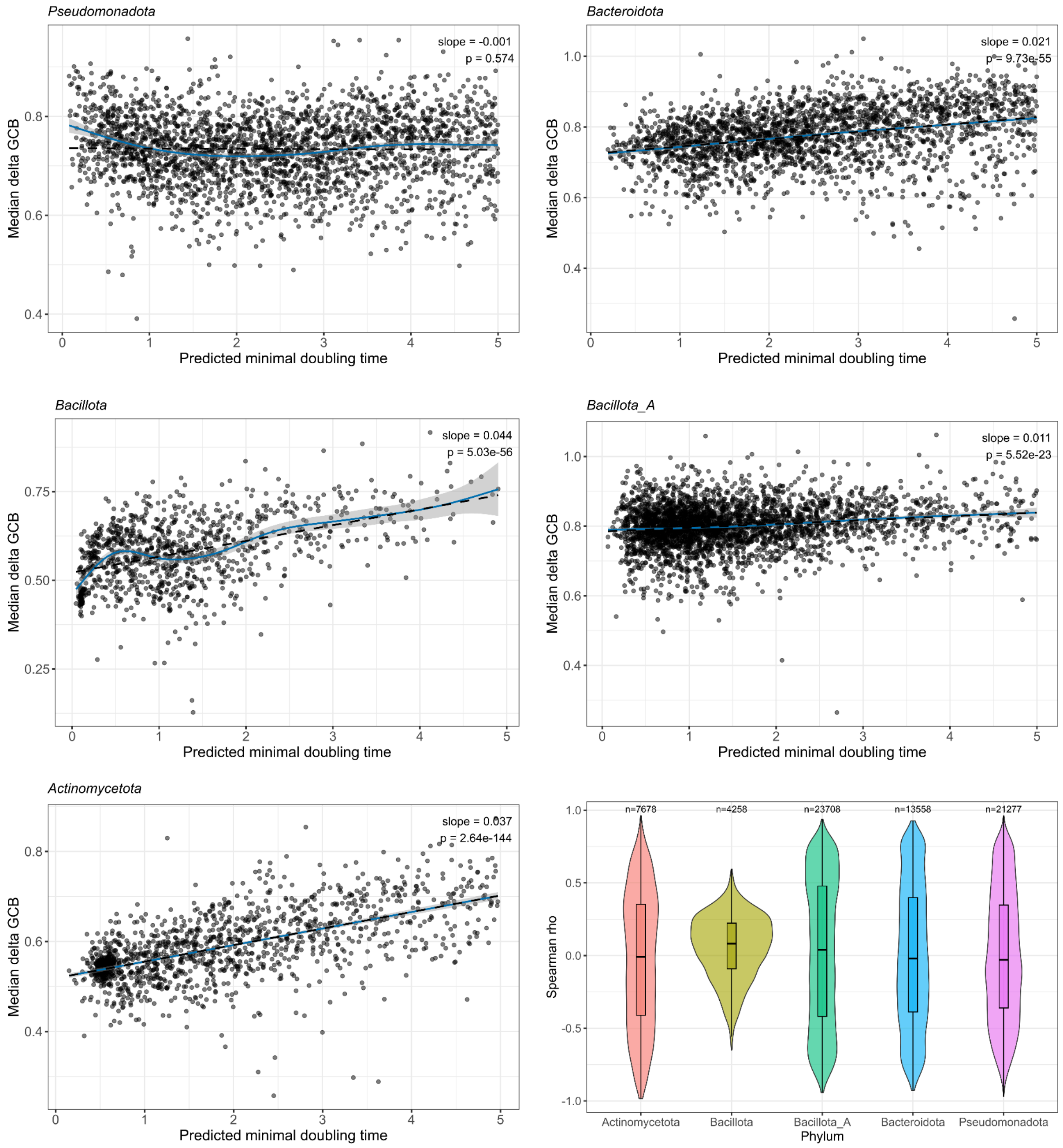


**Supplementary Fig. 15**

**Median |ΔGCBz| values slightly higher in slow compared to fast growing bacteria.** Using significant gene pairs (FDR < 0.01) in all five main phyla (**a**-**e**), we calculated Spearman correlations between |ΔGCBz| values of pairs and predicted minimal doubling time (PMDT) estimated with gRodon R package v.2.5.2 (Weissman et al. 2021).

We then calculated Spearman correlations between |ΔGCBz| values of pairs and PMDT values for each gene pair using the PMDT of genomes where each pair was present. As the distribution of correlations are slightly positive, this evidence indicates the dependence between |ΔGCBz| values of pairs and the growth rate is not strong, consistent with our models detecting significant pairs that are present across genomes with different PMDT.

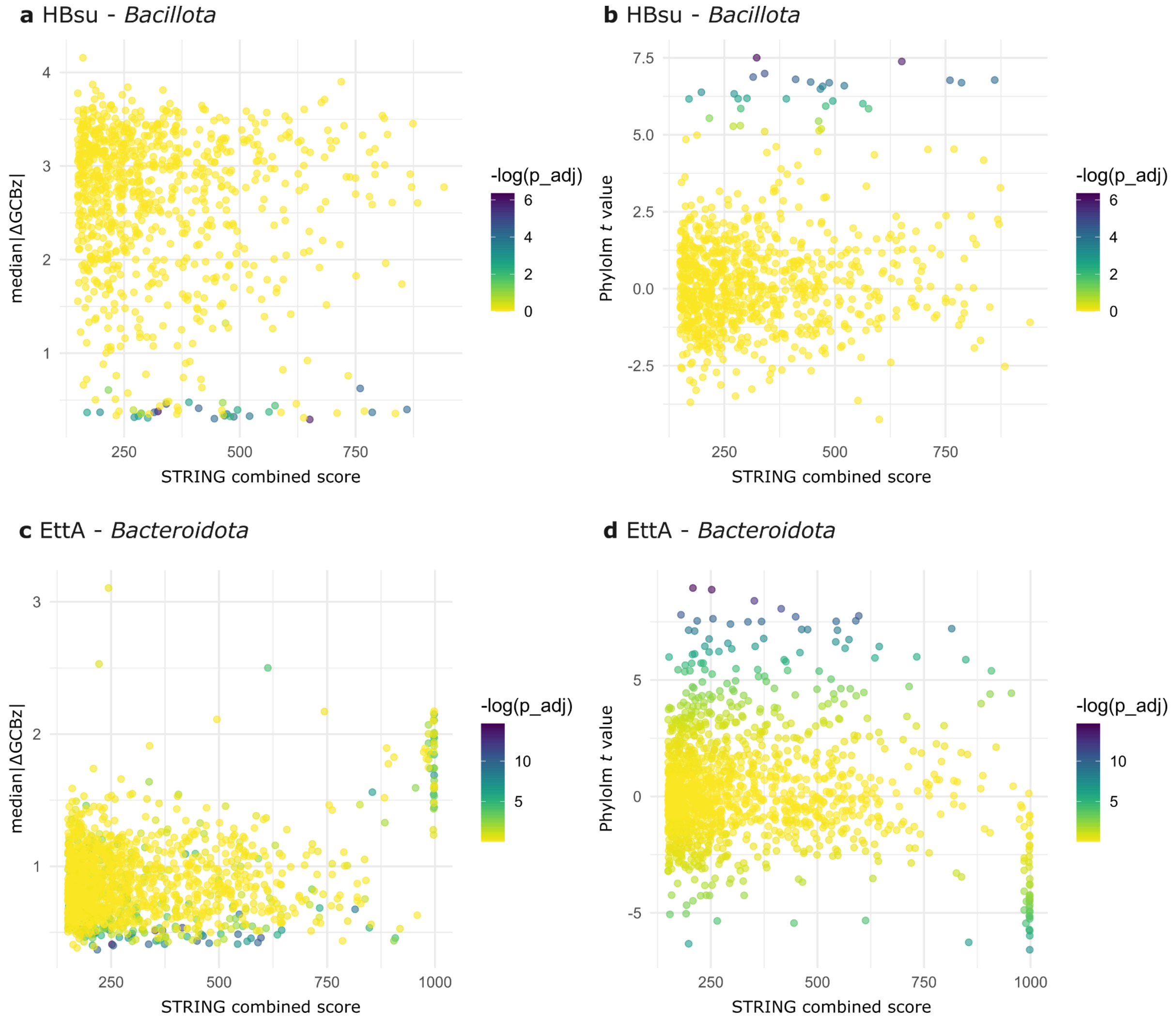


**Supplementary Fig. 16.**

**Median |ΔGCBz| and phylolm *t* values vs. STRING combined scores are shown for PGFam pairs corresponding to HBsu (PGF_01175502) in *Bacillota* and EttA (PGF_02059020) in *Bacteroidota*.** Genes corresponding to PGFam pairs across genomes were selected and their |ΔGCBz| was calculated per genome.

Median values across all genomes were calculated. A lower median |ΔGCBz| indicates genes have similar GCB values. Phylolm *t* values were obtained from SCUE-PGGI models for each PGFam pair.

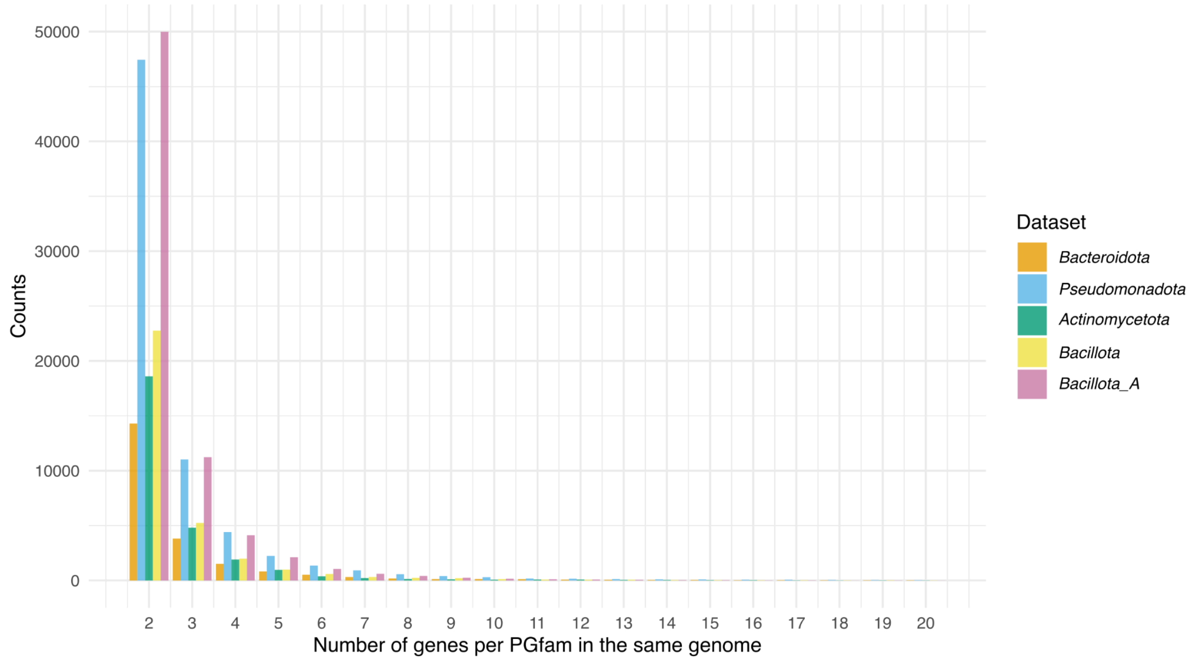


**Supplementary Fig. 17**

**Cases where PGFams have multiple representatives in the same genome.** Significant changes in CUB across PGFam trees were selected where the same genome had multiple representatives in the same PGFam, with at least one representative being significantly different compared to the others, and then classified according to the number of representatives in the same genome. Counts represent the number of cases across PGFams and genomes. Note that one PGFam may include multiple genomes with > 1 representative, and each genome may include multiple PGFams with > 1 representative.
